# Epigenetic Programming during thymic development sets the stage for optimal function in effector T cells via DNA demethylation

**DOI:** 10.1101/2021.05.03.442517

**Authors:** Athmane Teghanemt, Priyanjali Pulipati, Kenneth Day, Matt Yorek, Ren Yi, Kara Misel-Wuchter, Henry L Keen, Christy Au, Thorsten Maretzky, Prajwal Gurung, Dan R. Littman, Priya D. Issuree

## Abstract

The repressive effect of DNA methylation at promoters is well-known. However, its role within conserved sequences in intragenic and intergenic regions is less clear. Using *Cd4* as a model gene, here we show that DNA methylation regulates the function of stimulus-responsive regulatory elements in effector T cells. Two *cis*-elements orchestrate intra-and intergenic DNA demethylation of the *Cd4* gene during thymic development, which in turn licenses a stimulus-responsive element, E4a, for its later function in effector cells. Deficiency in DNA demethylation leads to impaired E4a function, reduced H3K4me3 promoter levels and an inability to repel *de novo* DNA methylation during replication, ultimately leading to gene silencing. This physiological reduction in CD4 expression leads to a defect in Th1 polarization during cutaneous Leishmaniasis. Similar patterns of regulation were observed in a broad number of genes, highlighting an essential role for DNA demethylation during thymic development in modulating the function of stimulus-responsive elements.

## INTRODUCTION

DNA methylation, specifically methylation of the fifth position of cytosine (5mC) in a CpG context, is a highly conserved and heritable epigenetic modification that modulates gene expression during mammalian development [1]. Compared to the well-documented negative correlation of 5mC at promoters and enhancers with gene expression, less is known about the function of 5mC at intragenic and intergenic regions. While in certain tissue types active genes exhibit more intragenic 5mC than repressed genes, this correlation appears to be tissue/cell-type specific, making the role of 5mC unclear [2–5] and necessitating the use of a tractable system to dissect the link between gene expression and intragenic methylation during cellular differentiation.

Thymic development of αβ CD4^+^ T cells provides a unique opportunity for the assessment of how DNA methylation influences stage-specific gene expression. CD4^+^ helper and regulatory T cells and CD8^+^ cytotoxic T cells originate from a common precursor, the double positive (CD4^+^CD8^+^ DP) thymocyte, and lineage specification is accompanied by dynamic changes in 5mC in many genes that continue to be expressed following exit from the thymus. It has been demonstrated that one of the first and longer-lasting intermediates of DNA demethylation, 5hmC, is abundant in newly-specified CD4^+^CD8^-^ thymocytes [6]. A large proportion of these 5hmC marks occur predominantly in intergenic and intragenic regions of active genes but are depleted at the transcription start sites (TSS) [6]. However, the roles of 5mC/5hmC at these sites in gene function remain to elucidated.

Using the *Cd4* locus as a tractable model gene in helper T cells, we previously showed that the locus contains extensive intragenic DNA methylation in the DP precursors, in which *Cd4* is expressed, as well as in the earlier CD4^-^CD8^-^ double-negative (DN) precursors, when it is not expressed [7], already suggesting that while intragenic methylation positively correlates with CD4 expression in DP precursors, it is negatively correlated in its earlier DN precursors. This underscores the importance of context. As DP precursors commit to the CD4^+^CD8^-^ lineages, the *Cd4* locus undergoes extensive TET1/3-mediated DNA demethylation [8]. We previously reported that this demethylation process is highly coordinated via stage-specific activity of *cis*-regulatory elements (CRE) at the locus. An upstream CRE (E4p), which dictates *Cd4* expression in DP precursors, and a downstream intronic CRE (E4m), which modulates *Cd4* expression during commitment and maturation of helper and regulatory T cells, are both required for TET1/3-mediated demethylation [8–10]. Lack of demethylation significantly reduced CD4 expression in activated proliferating T cells but did not compromise CD4 expression in naïve CD4^+^ T cells [8]. The mechanism of why expression is only dramatically affected in effector T cells remains unclear.

Herein, we discovered a novel stimulus-responsive CRE (E4a) that modulates *Cd4* expression in effector helper T cells. E4a is licensed during thymic T cell development and lack of DNA demethylation during development compromises its later function in effector cells. We found that reduced CRE function led to a breakdown in the maintenance of DNA demethylation during replication, consequently leading to promoter silencing of *Cd4*. We have interrogated the impact of reduced CD4 expression in effector T cells in a mouse model of cutaneous leishmaniasis and show that reduced CD4 expression during replication impairs Th1 differentiation, which correlates with defective clearance of infection. Furthermore, using a genome-wide analysis, we identified a subset of genes whose expression in effector T cells is highly dependent on proper epigenetic programming during thymic development. Together these findings demonstrate the importance of intragenic and intergenic demethylation during development in the modulation of stimulus-responsive CREs in effector T cells and whose function is required for preventing *de novo* methylation during replication.

## RESULTS

### E4a modulates *Cd4* expression in effector T cells in a partially redundant manner with E4m

We and others previously demonstrated a role for the E4m CRE in the upregulation of CD4 expression following positive selection in the thymus [8, 11]. Together with another CRE, E4p, which controls CD4 expression in DP thymocytes [9], E4m ensures sufficient CD4 expression for continuous and robust TCR signaling during lineage commitment, and is vital to ensure differentiation into helper and regulatory T cell lineages [8, 11]. E4m-deficient T cells in the periphery continued to display reduced CD4 expression, indicating that this enhancer function may be required to set the stage for optimal expression post-thymic maturation. Additionally, CD4 expression in *Cd4^E4mΔ/Δ^* T cells was further reduced in the course of T cell proliferation [8], indicating a possible role for E4m-dependent epigenetic regulation of *Cd4*. To first test the requirement of E4m in post-thymic T cells, we generated *Cd4^E4mflox/flox^* mice. Using a bicistronic Cre-recombinase GFP retroviral transduction approach [9], we induced recombination of E4m in *in vitro* activated CD4^+^ T cells. Unexpectedly, CD4 expression was not significantly affected in proliferating cells following deletion of E4m (**Fig. 1A, 1B, Supplementary Fig. 1A**) suggesting that E4m is dispensable for maintaining CD4 expression in activated T cells. We hypothesized that another CRE may compensate for maintaining CD4 expression in proliferating T cells in the absence of E4m. To test this, we used the Immgen ATAC-seq database to examine chromatin accessibility, as a readout for putative CREs in CD4^+^ thymic T cells, CD4 ^+^ naïve T cells and *in vitro* activated CD4^+^ T cells, and identified a region upstream of the *Cd4* TSS that was uniquely accessible in activated CD4^+^ T cells (**Fig. 1C**). We designated this putative CRE as E4a and generated *Cd4^E4aΔ/Δ^* and *Cd4^E4aΔ/Δ/ E4mΔ/Δ^* mice. CD4^+^ T cells from *Cd4^E4aΔ/Δ^* mice displayed a modest but significant reduction in CD4 expression upon activation (**Supplementary Fig. 1B,C**). However, combined loss of E4m and E4a resulted in an earlier and more profound loss of CD4 expression than loss of E4m alone (**Fig. 1D, 1E**). Strikingly, CD4 expression in *Cd4^E4aΔ/Δ/ E4mΔ/Δ^* T cells was markedly reduced following proliferation at 48h as compared to 16 h post T cell activation, during which time there is no cell division (**Supplementary Fig. 1D)**, suggesting a replication-coupled loss of expression. The profound reduction in CD4 expression in *Cd4^E4aΔ/Δ/ E4mΔ/Δ^* T cells was accompanied by reduced *Cd4* mRNA expression, consistent with compromised transcription (**Fig. 1F**). We further assessed whether loss of CD4 was due to alternately spliced *Cd4* transcripts, but found similar defects in mRNA levels when assessing usage of exons 1,2 and 3 (**Supplementary Fig. 1E, 1F**). Loss of CD4 expression in *Cd4^E4mΔ/Δ/ E4aΔ/Δ^* T cells was independent of the strength of TCR activation, as serial dilution of anti-CD3 stimulation led to equal loss of CD4 expression during proliferation (**Supplementary Fig. 1G, 1H**). Furthermore, when naïve CD4 helper T cells from *Cd4^E4mΔ/Δ/ E4aΔ/Δ^* were co-transferred with WT naïve CD45.1 helper T cells into *Rag^-/-^* mice (**Fig. 1G**), CD45.1 WT T cells maintained high CD4 levels during homeostatic proliferation while *Cd4^E4aΔ/Δ/ E4mΔ/Δ^* T cells showed a dramatic downregulation (**Fig. 1H, 1I**). Together our data suggest that E4a is a stimulus-responsive CRE and acts in a partially redundant manner with E4m to maintain CD4 expression in proliferating CD4^+^ T cells.

**Fig. 1:**
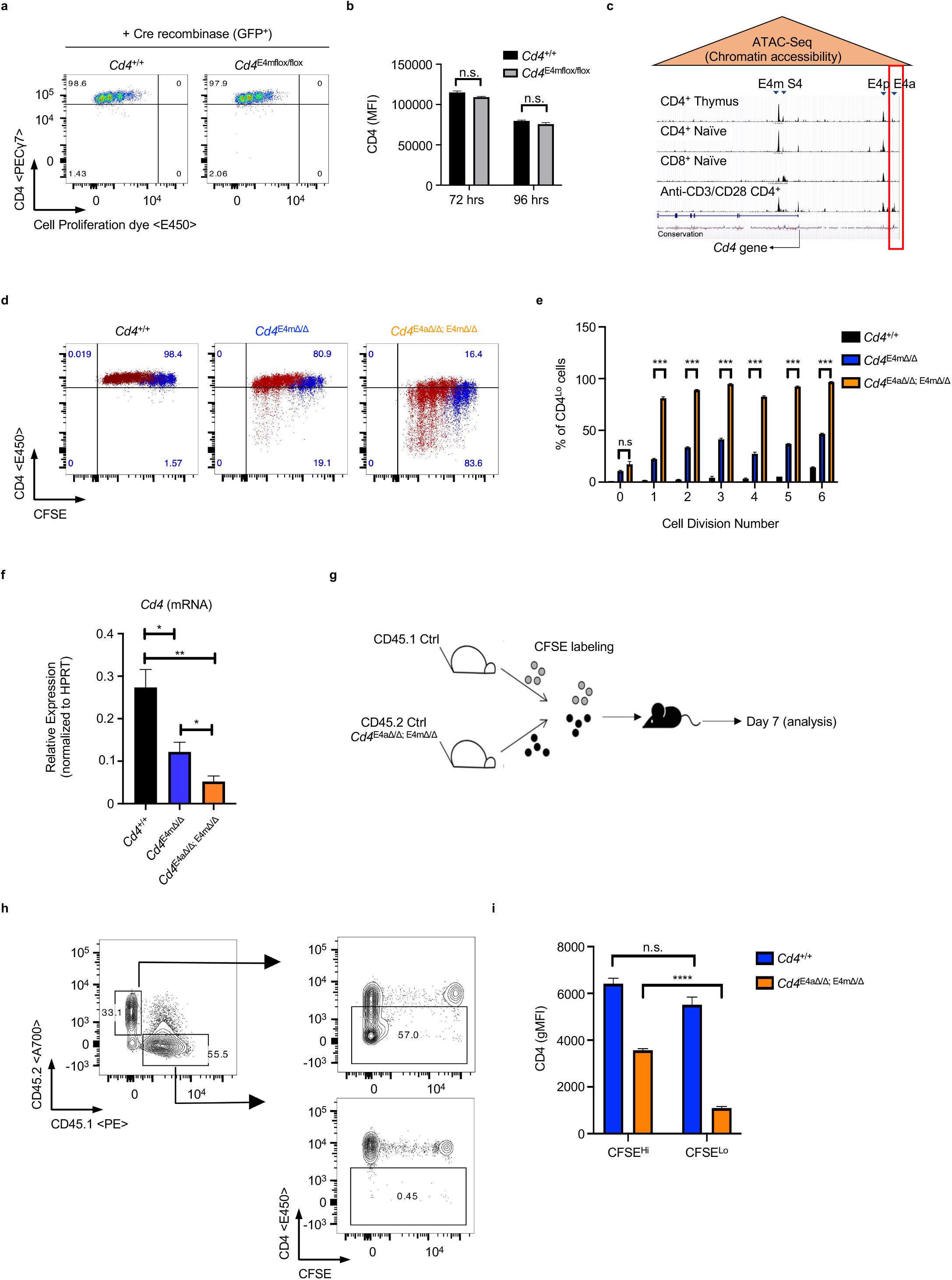
E4a modulates *Cd4* expression in effector T cells in a partially redundant manner with E4m. **a,** Flow cytometry plot of CD4 expression in control and *Cd4^E4mflox/flox^* T cells, transduced with Cre-recombinase and labeled with a cell proliferation dye prior to activation. Transduced cells were gated on GFP. Data is representative to 3 independent experiments. **b**, CD4 MFI of cells from (a) measured at two different time-points post transduction. Data shown is representative of 3 experiments and expressed as mean ± SD of 3 technical replicates. **c**, Integrative Genome View (IGV) browser shot of ATAC-Seq data depicting chromatin accessibility within and upstream of the *Cd4* locus. Tracks shown are set at the same scale. Arrow indicate the annotated *Cd4* TSS and red box denotes the location of E4a. **d**, FACS plot showing CD4 expression in proliferating T cells that were CFSE-labeled prior to *in vitro* activation with anti-CD3/CD28. Dot plots at 72hrs (Blue dots) and 96hrs (Red dots) post-activation were overlaid. Data is representative of >3 independent experiments. **e**, Percent of cells with low CD4 expression at indicated proliferation cycles assessed by CFSE labeling. Data shown is representative > 3 experiments and expressed as mean ± SD of 3 technical replicates. ***p<0.001 (Student t test). **f**, CD4 mRNA expression (exon 3-4) in activated CD4 T cells. RNA was isolated 96hrs post activation with anti-CD3/CD28. Data is expressed as mean ± SEM of 4-5 individual mice/group. *p<0.05, **p<0.01 (One-Way ANOVA and Bonferroni test). **g**, Schematic illustration of experimental design. Naïve WT CD45.1 T cells or CD45.2 T cells from *Cd4^E4mΔ/Δ/ E4aΔ/Δ^* mice were FACS-sorted, CFSE labeled and mixed at a 1:1 ratio before i.v. transfer into *Rag^-/-^* recipients. 7 days later, LNs and spleen were isolated and analyzed by Flow cytometry. **h**,FACS plot showing CD4 expression in CD4 T cells from *Rag^-/-^* mice at day 7 post transfer. Cells were gated on congenic markers to distinguish WT and mutant T cells. Data is representative of 2 independent experiments with 3-5 mice/experiment. **i**, CD4 geometric MFI in cells gated on CSFE staining profiles. Data is expressed as mean ± SEM of 3-5 individual mice/group and is representative of 2 independent experiments. **** p<0.0001 (unpaired Student’s t test)

### E4a is a stimulus-responsive *cis*-regulatory element licensed during development

We next assessed whether E4a was required during T cell development. Deletion of E4a alone had no effect on CD4 expression in pre-selected TCRβ^lo^CD24^hi^CD69^−^ DP thymocytes, nor in recently selected CD4^+^TCRβ^hi^CD24^hi^CD69^+^, mature CD4^+^TCRβ^hi^CD24^lo^CD69^−^ thymocytes or naïve peripheral CD4^+^ T cells (**Supplementary Fig. 2A, 2B**). *Cd4^E4aΔ/Δ/ E4mΔ/Δ^* mice mirrored *Cd4^E4mΔ/Δ^* mice with normal CD4 levels in pre-selected DP thymocytes and reduced expression in recently selected and mature CD4^+^ thymocytes compared to *Cd4^+/+^* controls (**Fig. 2A, 2B**). In peripheral CD4 T cells, *Cd4^E4mΔ/Δ/ E4aΔ/Δ^* mice displayed a modest but statistically significant reduction in CD4 expression compared to *Cd4^E4mΔ/Δ^* mice (**Fig. 2C, 2D, Supplementary Fig. 2C, 2D**). These results suggest that E4a modulates CD4 expression in peripheral and proliferating CD4^+^ T cells (**Fig. 1D, Supplementary Fig. 1C**). In support of this interpretation, ChIP-Seq analysis revealed the presence of both H3K27Ac and p300, which mark enhancer-like regions [12], within the E4a region in activated CD4^+^ T cells (**Fig. 2F**) but not in DP^+^ and CD4^+^ thymocytes (**Fig. 2G**). In contrast, H3K27Ac marks were highly enriched at the E4p and E4m regions in DP^+^ and CD4^+^ thymocytes, respectively (**Fig. 2G**), in agreement with their established enhancer activities during development [8, 9]. Notably, there was enrichment of H3K4me1 marks, which demarcate primed and poised enhancers [12], at the E4a region in DP^+^ and CD4^+^ thymocytes but not in activated CD4^+^ T cells (**Fig. 2F, 2G**), suggesting that while E4a is not active and not required during development of CD4-lineage T cells, it is primed early during thymic differentiation.

**Fig. 2:**
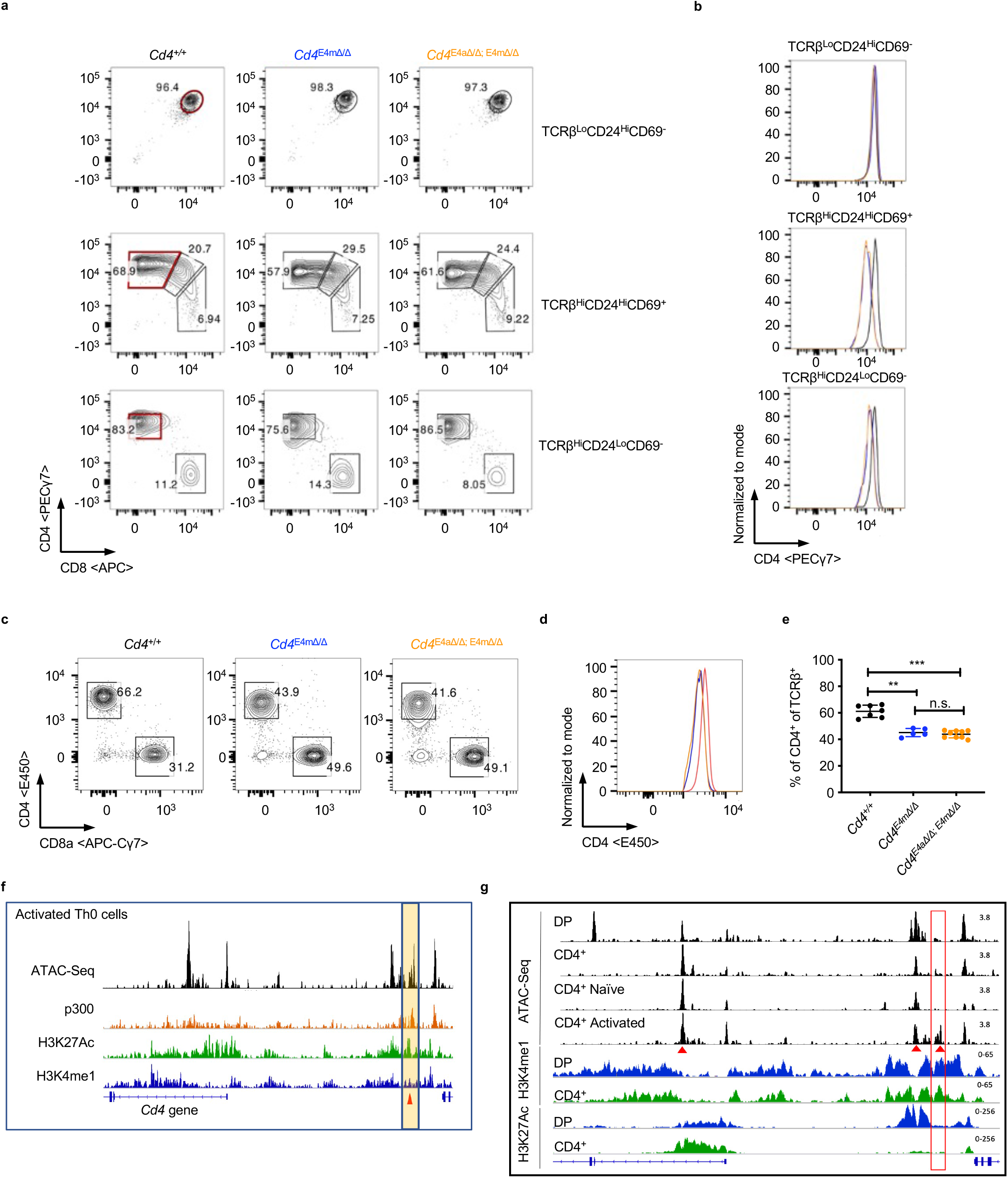
E4a is a stimulus-responsive *cis*-regulatory element licensed during development. **a,b,c**, Flow cytometry analysis of T lymphocytes from *Cd4^+/+^, Cd4^E4mΔ/Δ^* and *Cd4^E4mΔ/Δ/ E4aΔ/Δ^* mice. **a**, FACS contour plots showing pre-selected TCRβ^lo^CD69^-^CD24^hi^DP (top panel), recently selected TCRβ^hi^CD69^+^CD24^hi^ (middle panel) and mature TCRβ^hi^CD69^-^CD24^lo^ T cell populations (bottom panel) in the thymus. **b**, Histogram overlay of CD4 expression gated on CD4 populations at different stages in maturation denoted in red boxes in (a). **c**, FACS contour plots showing CD4 and CD8 proportions among TCRβ+ T cells from the spleen/LN. **d**, Histogram overlay of CD4 expression gated on TCRβ+ CD4^+^ T cells from the spleen/LN of mice. Data is representative of >3 experiments with 5 animals/group/experiment. **e**, Frequency of CD4^+^ T cells among TCRβ+ T cells in the spleen/LN of mice with indicated genotypes. Data are expressed as mean ± SEM of individual mice (n = 5 of two independent experiments). **p<0.01, ***p<0.001 (One-Way ANOVA and Bonferroni test). **f**, IGV snapshots of the *Cd4* locus in activated T (Th0) cells depicting accessibility sites by ATAC-Seq, p300 binding and presence of H3K27Ac and H3K4me1 marks by ChIP-Seq. Highlighted yellow box and arrow indicates the location of E4a. **g**, IGV snapshots of the *Cd4* locus depicting accessibility sites by ATAC-Seq in thymocytes and activated T cells, and H3K27Ac and H3K4me1 signatures by ChIP-Seq in DP and CD4+ thymocytes. Red box indicates the location of E4a and red arrow denotes the location of E4m, E4p and E4a.

### Lack of DNA demethylation during development affects the function of E4m/E4a in effector CD4-lineage T cells

The role of intragenic methylation in the regulation of *Cd4* gene expression remains paradoxical. Tet1/3 ^cDKO^ naïve helper T cells (*Rorc(t)Cre*^Tg^ *Tet1/3^fl/fl^*) have similar expression of CD4 compared to WT naïve T cells despite significantly higher intragenic DNA methylation [8]. However, Tet1/3^cDKO^ helper T cells have substantial loss of CD4 expression upon proliferation, similarly to *Cd4^E4mΔ/Δ^* mice [8]. We thus hypothesized that DNA methylation compromises the function of E4m/E4a in effector CD4-lineage T cells, potentially explaining the need for demethylation during development. Notably, there are no CpG motifs in the E4a CRE. However, DNA methylation marks, determined by CATCH-Seq as previously described [7, 8, 13], were present in regions flanking the E4a site as well as the E4m region in naïve Tet1/3 cDKO compared to naïve WT controls, (**Fig. 3A, Supplementary Fig. 3A**, **3B**). Furthermore, we found a significant reduction in H3K4me3 at the promoter region of *Cd4* (**Fig. 3B**) and a significant gain of H3K9me3 at the E4a region in Tet1/3^cDKO^ naïve helper T cells compared to controls (**Fig. 3C**), suggestive of a less permissive chromatin environment. Activated Tet1/3^cDKO^ helper T cells also displayed reduced H3K4me3 at the *Cd4* promoter and elevated H3K9me3 at the E4a region, concordant with reduced function of E4a during T cell proliferation (**Supplementary Fig. 3C, 3D**). Since the defect in CD4 expression was less profound in Tet1/3^cDKO^ compared to *Cd4^E4mΔ/Δ/ E4aΔ/Δ^* effector T cells (**Supplementary Fig. 3E**), we reasoned that all enhancer activity was lost in *Cd4^E4mΔ/Δ/ E4aΔ/Δ^* effector T cells whereas enhancer function was only partially impaired in Tet1/3^cDKO^ T cells. To test this, T cells were treated with an inhibitor to the histone acetyltransferases p300/ cAMP response element binding protein (CBP) which are key transcriptional activators recruited to enhancers and important for their function [14–16]. In agreement with our hypothesis, inhibition of p300/CBP activity reduced CD4 expression in a dose-dependent manner in control and Tet1/3^cDKO^ T cells but had no impact on *Cd4^E4mΔ/Δ/ E4aΔ/Δ^* T cells (**Supplementary Fig. 3E**). Next, we examined DNA methylation at the *Cd4* locus in *Cd4^E4aΔ/Δ^* and *Cd4^E4mΔ/Δ/ E4aΔ/Δ^* activated effector T cells. E4a deletion resulted in modestly increased DNA methylation downstream of the *Cd4* TSS and in the upstream intergenic region (**Fig. 3D, 3E**). In contrast, activated *Cd4^E4mΔ/Δ/ E4aΔ/Δ^* CD4 T cells displayed hyper DNA methylation both in the *Cd4* promoter and downstream of the TSS (**Fig. 3E**), which correlated with the dramatic loss of CD4 in these cells. Taken together, we surmise that DNA demethylation during thymic development plays an important role in shaping the epigenetic landscape for optimal regulation of *Cd4* expression in effector CD4-lineage T cells.

**Fig. 3:**
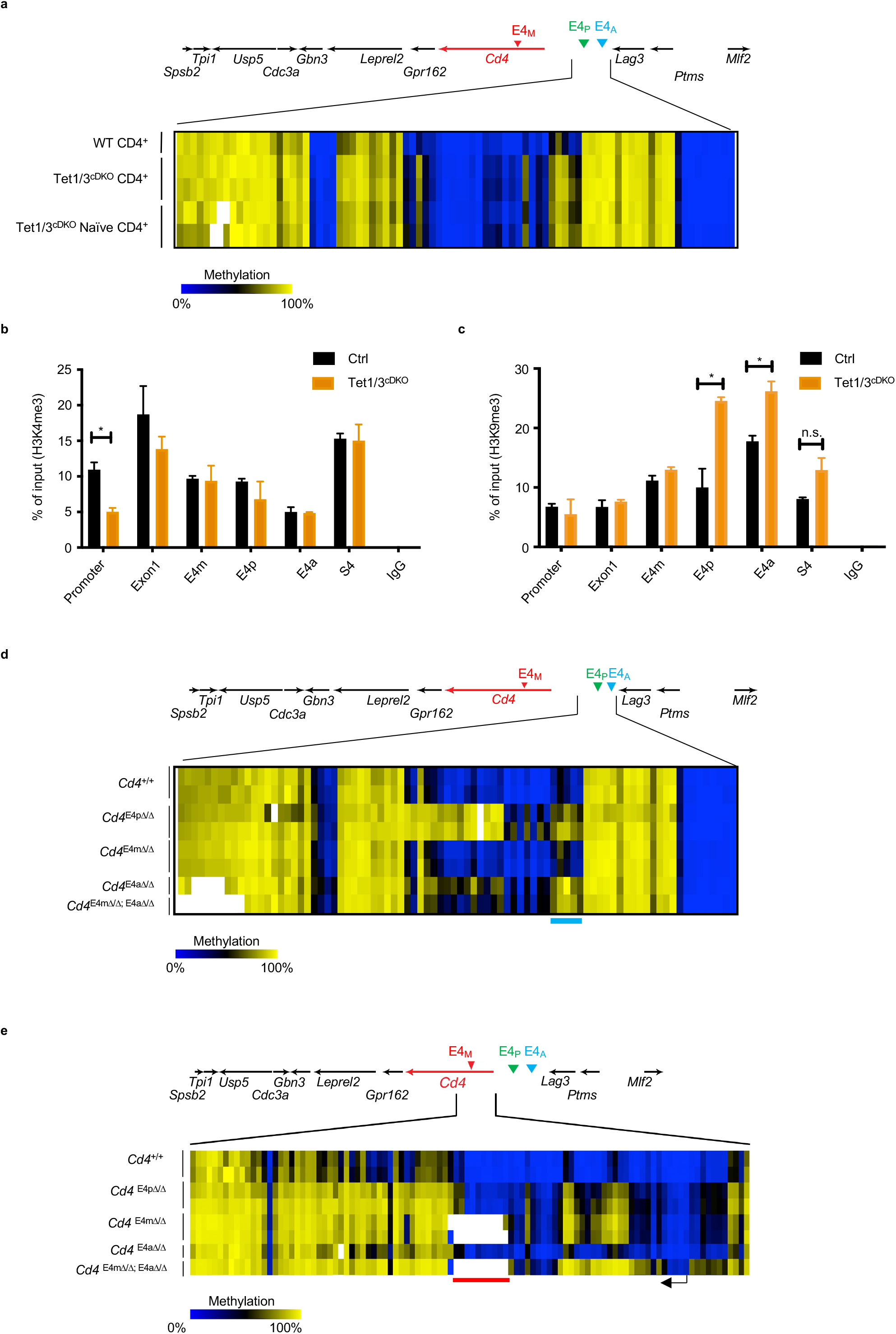
Lack of DNA demethylation during development affects the function of E4m/E4a in effector CD4-lineage T cells. **a**, Heatmap depicting percent CpG methylation in Control CD4+ (Tet1/3^flox/flox^), Tet1/3^cDKO^ CD4+ mature thymocytes and Tet1/3^cDKO^ CD4+ naïve peripheral T cells for CpGs -9270bp to -15869bp relative to the *Cd4* TSS (Chr6:124847307-124853906; mm9). The blue line underlines CpGs flanking the E4a region. CATCH-seq was performed on genomic DNA from sorted populations of TCRβ^hi^CD24^lo^CD69^−^CD4^+^CD8^−^ thymocytes or CD4^+^TCRβ^+^CD62L^hi^ CD44^-^ T cells from LN/Spleen. Replicates are from 2 independent mice. **b**, H3K4me3 modifications assessed by ChIP-qPCR in sorted naïve CD4 T cells. Data are averaged from two biological replicates/genotype and representative of 2 experiments. *p<0.05 (Student’s t test) **c**, H3K9me3 modifications assessed by ChIP-qPCR in sorted naïve CD4 T cells. Data are averaged from two biological replicates/genotype and representative of 2 experiments. *p<0.05 (Student’s t test). **d**, Heatmap depicting percent CpG methylation in activated (WT) *Cd4^+/+^, Cd4^E4pΔ/Δ^*, *Cd4^E4mΔ/Δ^*,*Cd4^E4aΔ/Δ^* and *Cd4^E4mΔ/Δ/ E4aΔ/Δ^* T cells for CpGs -9270bp to -15869bp relative to the *Cd4* TSS (Chr6:124847307-124853906; mm9). The blue line underlines CpGs flanking the E4a region. CATCH-seq was performed on genomic DNA from naïve T cells activated i*n vitro* with anti-CD3/CD28 for 120hrs. Note data for *Cd4^E4pΔ/Δ^* and *Cd4^E4mΔ/Δ^* conditions were from previously published experiments with similar experimental conditions [7, 8]. **e**, Heatmap depicting percent CpG methylation in WT *Cd4^+/+^, Cd4^E4pΔ/Δ^*, *Cd4^E4mΔ/Δ^*,*Cd4^E4aΔ/Δ^* and *Cd4^E4mΔ/Δ/ E4aΔ/Δ^* T cells for CpGs from +6200 to −669 relative to the *Cd4* TSS (Chr6:124832027–124838896; mm9). A red line underlines CpGs in E4m (indicated by the gap in the mutant mice) and a black arrow indicates the *Cd4* TSS. Red box indicates the *Cd4* promoter. Note data for *Cd4^E4pΔ/Δ^* and *Cd4^E4mΔ/Δ^* conditions were from previously published experiments with similar experimental conditions [7, 8].

### Reduced enhancer activity as a result of DNA methylation leads to *Cd4* promoter silencing during replication of effector CD4^+^ T cells

We next postulated that hypermethylation of the *Cd4* promoter in activated *Cd4^E4mΔ/Δ/ E4aΔ/Δ^* T cells is the cause of loss of *Cd4* expression and is a consequence of the reduction in transcriptional activity directed by E4m and E4a. We therefore tested whether boosting enhancer activity prevents DNA methylation at the promoter and restores *Cd4* expression. To do so, we took advantage of the transcription factor TCF1, which we previously demonstrated binds to the CRE E4p [8]. We had shown that overexpression of β-catenin, a co-activator of TCF-1, in activated *Cd4^E4mΔ/Δ/^* T cells fully restored CD4 expression in the absence of E4m by maintaining E4p-mediated transcription [8] (**Fig. 4A**). However, overexpression of β-catenin in *Cd4^E4mΔ/Δ/ E4aΔ/Δ^* T cells led to only a partial rescue of CD4 expression (**Fig. 4B, 4C**), while CD4 expression was restored to control levels in in *Cd4^E4mΔ/Δ^* and Tet1/3^cDKO^ T cells (**Fig. 4C, 4D**), suggesting that E4p activity is not sufficient to rescue CD4 expression in effector cells lacking both E4m and E4a. To test whether rescue of CD4 expression in *Cd4^E4mΔ/Δ^* T cells by TCF1/β-catenin was a consequence of changes in DNA methylation, we examined *Cd4^E4mΔ/Δ^* T cells that lost CD4 expression during replication (CD4^Lo^) and β-catenin-transduced *Cd4^E4mΔ/Δ^* CD4 T whereby CD4 expression was fully restored (CD4^Hi^). CD4^Hi^ *Cd4^E4mΔ/Δ^* T cells displayed a remarkable absence of DNA methylation proximal to the *Cd4* TSS, while intragenic methylation was unchanged (**Fig. 4E**), indicating that β-catenin most likely prevents the establishment of *de novo* methylation at the promoter, which causes a loss of CD4 expression. These results also suggest that methylation in the intragenic region of *Cd4* does not directly impede gene expression as it was unchanged when CD4 expression was restored.

**Fig. 4:**
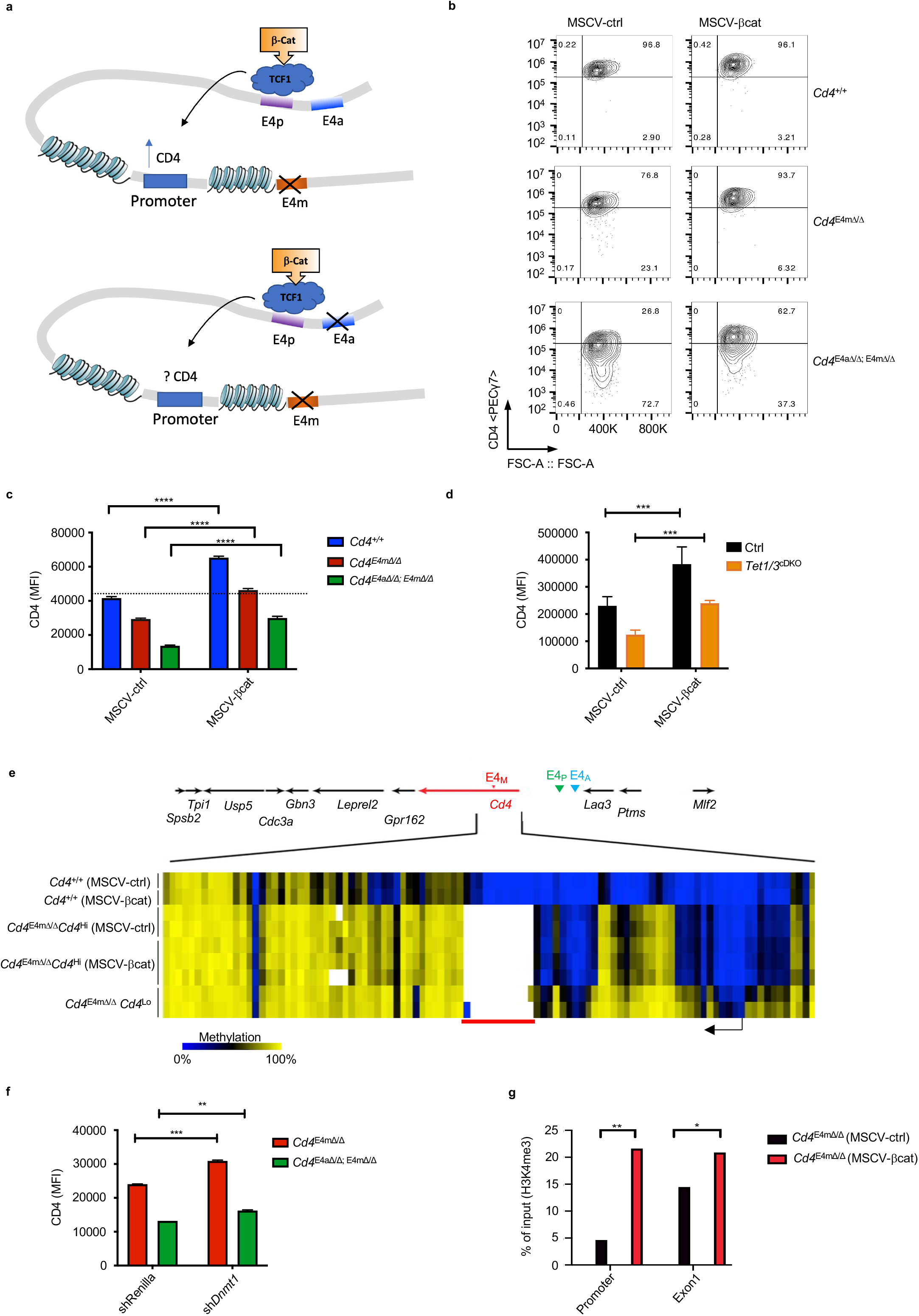
Reduced enhancer activity as a result of DNA methylation leads to *Cd4* promoter silencing during replication of effector CD4^+^ T cells. **a**, Schematic illustration depicting the location and orientation of the *Cd4* enhancers, and TCF1 binding site previously validated [8]. **b,** FACS Contour plots showing expression of CD4 in activated T cells with indicated genotypes upon rescue with a control vector or a vector overexpressing β-Catenin. Data shown are 96hrs post retroviral transduction and representative of >3 independent experiments. **c,** CD4 gMFI on T cells analyzed 96hrs post retroviral transduction with a control vector or β-Catenin vector. Data shown is representative of 3 experiments and expressed as mean ± SD of 3 technical replicates. ****p<0.0001 (2-way ANOVA and Sidak’s test)**. d,** CD4 gMFI on T cells analyzed 96hrs post retroviral transduction with a control vector or β-Catenin vector. Data shown is representative of 3 experiments and expressed as mean ± SD of 3 technical replicates. ***p<0.001(One-way ANOVA and Bonferroni test)**. e,** Heatmap depicting percent CpG methylation CpGs from +6200 to −669 relative to the *Cd4* TSS (Chr6:124832027–124838896; mm9) in activated WT CD4^+^ and *Cd4^E4mΔ/Δ^* CD4+ T cells transduced with a control vector or β-Catenin. A red line underlines CpGs in E4m (indicated by the gap in the mutant mice) and a black arrow indicates the *Cd4* TSS. CATCH-seq was performed on genomic DNA from *Cd4^E4mΔ/Δ^* GFP+ sorted populations (control vector or β-Catenin transduced). Data from replicates are from >/ 2 independent mice. Note data for CD4^Lo^*Cd4^E4mΔ/Δ^* and *Cd4^+/+^* conditions were from previously published experiments [8]. **f,** CD4 gMFI of *Cd4^E4mΔ/Δ^* and *Cd4^E4mΔ/Δ/ E4aΔ/Δ^* CD4^+^ T cells transduced with a shRNA against Renilla or Dnmt1. Cells were analyzed 96hrs post transduction and gated on GFP. Data shown is representative of 3 experiments and expressed as mean ± SD of 3 technical replicates. **p<0.01, ***p<0.001 (One-way ANOVA and Bonferroni test). **g**, H3K4me3 modifications assessed by ChIP-qPCR in control or β-Catenin transduced (GFP+) CD4^+^ T cells from *Cd4^E4mΔ/Δ^* mice. Cells were FACS sorted 96hrs post transduction. Data is representative of 2 independent experiments. ** p<0.01 (Student t test)

We next assessed whether gain of methylation at the *Cd4* promoter during replication was due to activity of the maintenance methyltransferase DNMT1 and/or the *de novo* methyltransferase DNMT3a [1]. To exclude a confounding role on gene regulation by the silencing element S4, which mediates suppression of CD4 in cytotoxic CD8 T cells [17–19], *Rorc(t)Cre*^Tg^ *Cd4*^S4 flox/flox^ mice (which lack both E4m and S4)[10, 19] were bred to *Dnmt3a* ^flox/flox^ mice. In the absence of DNMT3a, we observed a significantly lower proportion of T cells with reduced CD4 expression during replication as well as an increase in expression of CD4 compared to controls (**Supplementary Fig. 4A, 4B, 4C**). In addition, shRNA knockdown of *Dnmt1* in *Rorc(t)Cre*^Tg^ *Cd4*^S4 flox/flox^; *Dnmt3a^flox/flox^* T cells led to further rescue in CD4 expression (**Supplementary Fig. 4D**), suggesting that both DNMT1 and DNMT3a have roles in *Cd4* silencing during replication. Knockdown of *Dnmt1* in *Cd4^E4mΔ/Δ/ E4aΔ/Δ^* T cells led to a modest rescue of CD4 expression (**Fig. 4F**), supporting a partial contribution of DNA methylation to the silencing of *Cd4* in the absence of E4m and E4a, although it is noteworthy that only a very modest rescue was achievable in activated *Cd4^E4mΔ/Δ/ E4aΔ/Δ^* T cells due to the timing of shRNA knockdown. To elucidate how β-catenin prevented *de novo* methylation, we next assessed H3K4me3 levels. Compared to control vector-transduced T cells, β-catenin-transduced *Cd4^E4mΔ/Δ^* T cells had significantly higher enrichment of H3K4me3 marks at the promoter and TSS sites (**Fig. 4G**), suggesting that H3K4me3 prevents the gain of new DNA methylation, in support of the antagonistic relationship between H3K4me3 and DNA methylation [20, 21]. Together these findings suggest that a lack of thymic DNA demethylation impairs the activity of E4a and E4m in effector T cells, which is required to maintain high H3K4me3 at the *Cd4* promoter during activation and replication, and repel *de novo* DNA methylation that leads to *Cd4* gene silencing (**Supplementary Fig. 4E**).

### Reduced CD4 expression in effector T cells impairs parasitic clearance during Leishmaniasis

The need for continuous regulation of the *Cd4* gene in proliferating T cells brought into focus the importance of CD4 expression in effector T cells. Indeed, while CD4 coreceptor levels can vary in T-cell subsets *in vitro* [22], whether differential CD4 levels has a role in dictating cell fate of effector CD4 T cells *in vivo* is unclear. However, TCR signaling has been implicated in dictating the fate of differentiating effector T cells during infection [23, 24]. We therefore tested the biological impact of reduced CD4 expression in a *Leishmania major* (*L.major*) infection model which induces a canonical Th1 response in WT C57BL/6 mice [25]. Footpads of *Cd4^+/+^* and *Cd4^E4mΔ/Δ/ E4aΔ/Δ^* mice were cutaneously infected with *L. major* and swelling was assessed over time. Interestingly, while all animals developed footpad swelling, *Cd4^E4mΔ/Δ/ E4aΔ/Δ^* mice displayed delayed swelling relative to *Cd4^+/+^* mice. However, inflamed lesions ultimately resolved at the same time in *Cd4^+/+^* and *Cd4^E4mΔ/Δ/ E4aΔ/Δ^* mice (**Fig. 5A**). In contrast, no difference in footpad swelling was seen in *Cd4^E4mΔ/Δ^* or *Cd ^E4aΔ/Δ^* mice relative to *Cd4^+/+^* mice (**Fig. 5B, 5C**). However, *Cd4^E4mΔ/Δ/ E4aΔ/Δ^* mice showed significantly elevated parasitic burden in the footpad relative to controls both at the peak of infection (d28) and at the resolution of inflammation (d54) but not early in infection (d9), suggesting that delayed inflammatory swelling in *Cd4^E4mΔ/Δ/ E4aΔ/Δ^* correlates with impaired parasitic clearance (**Fig. 5D, 5E, 5F**). Activated CD44^hi^ T cells from the draining lymph nodes (dLNs) of infected *Cd4^E4mΔ/Δ/ E4aΔ/Δ^* mice displayed a dramatic downregulation of CD4 relative to *Cd4^+/+^* or *Cd4^E4mΔ/Δ^* and *Cd ^E4aΔ/Δ^* mice at day 9 (**Fig. 5G, 5H, Supplementary Fig. 5A, 5B, 5C**). Furthermore, the percentage of cells with reduced CD4 expression in *Cd4^E4mΔ/Δ/ E4aΔ/Δ^* mice significantly increased over time (**Fig. 5I**), recapitulating the phenotypic loss of CD4 expression upon proliferation seen *in vitro* and in *Rag^-/-^* mice (**Fig. 1D,1H**). Together these findings suggest that compromised TCR signaling due to unstable CD4 expression in helper T cells during *L. major* infection results in delayed inflammation and defective parasite clearance.

**Fig. 5:**
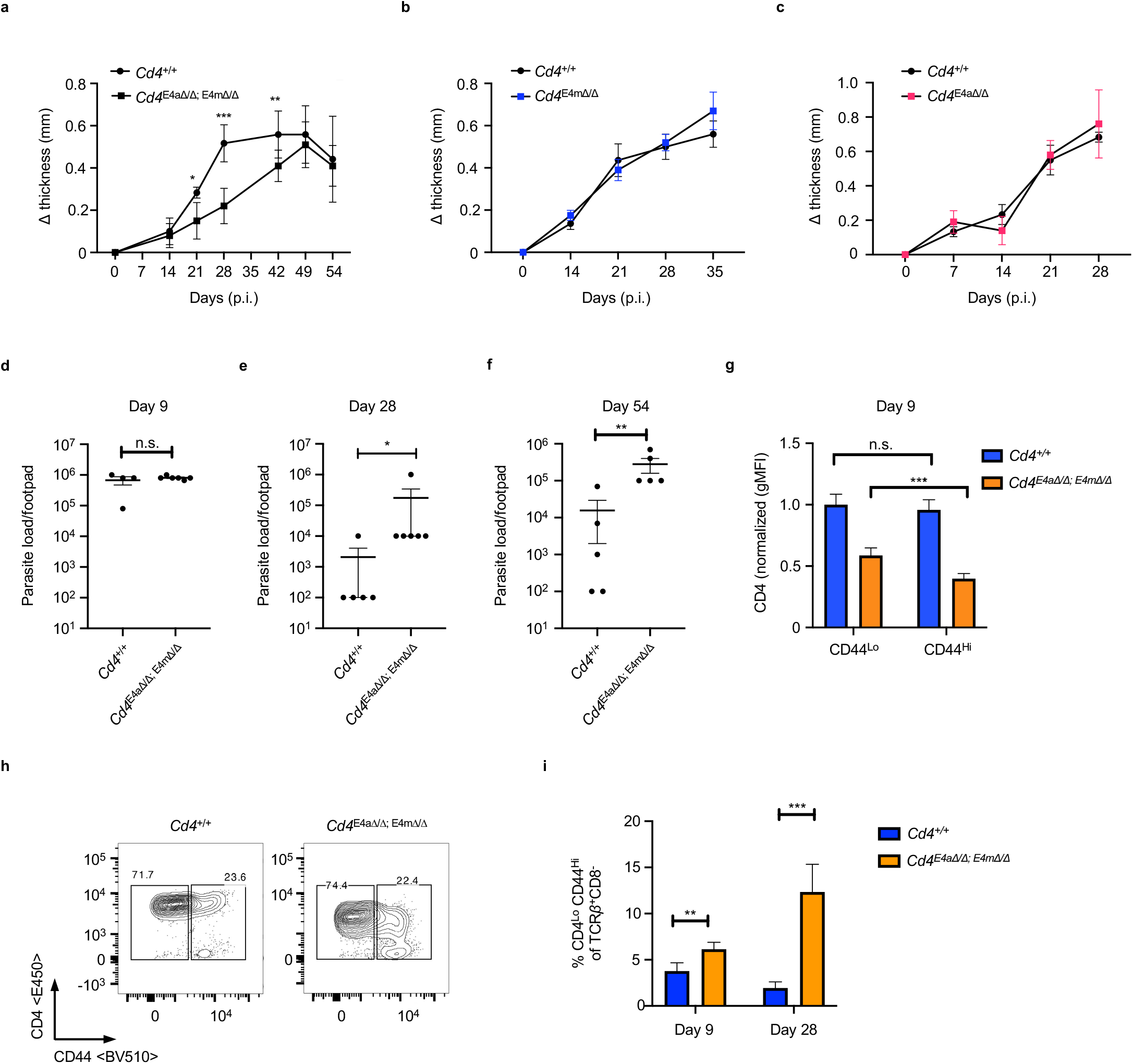
Reduced CD4 expression in effector CD4 T cells impairs parasitic clearance during Leishmaniasis. **a-c**, Footpads of indicated genotypes were infected with 2x 10^6^ *L. major* promastigotes. Footpad swelling was measured over days post infection (p.i). Data shown is mean ± SEM of indicated mice/group. *p<0.05, ** p<0.01, *** p<0.001 (Mann-Whitney test). **d-f**, Parasitic burdens in the footpads of infected mice. On the indicated days p.i., mice were euthanized, and *L. major* titers in the footpads were determined using a limiting dilution assay as described in Methods. Data shown is mean ± SEM of 4-5/group and representative of 2-4 independent experiments *p<0.05, ** p<0.01, *** p<0.001 (Mann-Whitney test). **g**, Normalized CD4 gMFI expression on CD8^-^ TCRβ^+^ CD44^Lo^ or CD8^-^ TCRβ^+^ CD44^Hi^ T cells from the inguinal dLNs of infected mice 9 days p.i. Data shown is a mean ± SEM of 9-10 mice/genotype and is a summary of 2 independent experiments. **h**, FACS contour plot showing CD4 versus CD44 expression among CD8^-^ TCRβ^+^ T cells from the inguinal dLNs of infected mice 9 days p.i. Data shown is representative of 2 independent experiments with 4-5 mice/group/experiment. **i**, Proportions of helper T cells with low CD4 expression among CD8^-^ TCRβ^+^ CD44^Hi^ T cells from the inguinal dLNs of infected mice, 9 and 28 days p.i. Data shown is mean ± SEM of 9-10 mice/group and is a summary of 2 or more independent experiments.

### Reduced CD4 expression impairs the differentiation of Th1 cells during Leishmaniasis

We next examined whether loss of CD4 impacts the antigen-specific response to *L.major* by assessing the proportion of CD11a^hi^CD49d^+^ and CD11a^hi^CD44^+^ T cells in the dLNs of *L.major*-infected mice. CD11a in combination with CD49d or CD44 are routinely used as surrogate markers to track antigen-experienced CD4^+^ T cells in viral and parasitic infections [26–30]. At day 9 p.i, the proportion of both CD11a^hi^CD49d^+^ and CD11a^hi^CD44^+^ T cells in the dLNs of *L.major* infected mice was significantly higher than in uninfected controls (**Fig. 6A, 6B**). No significant differences were observed in the proportion of these cells in CD4^+/+^ compared to *Cd4^E4mΔ/Δ/ E4aΔ/Δ^* mice, suggesting that reduced CD4 expression did not affect the ability of T cells to engage TCR signaling. However, antigen-experienced *Cd4^E4mΔ/Δ/ E4aΔ/Δ^* T cells had lower expression of CD5 and Nur77, suggestive of reduced TCR signal strength (**Fig. 6C, 6D**) [31, 32]. At 28 days post infection (d.p.i.), *Cd4^E4mΔ/Δ/ E4aΔ/Δ^* mice had a significantly lower frequency of Tbet^+^CD11a^hi^CD44^+^ helper T cells in the dLNs compared to CD4^+/+^ controls (**Fig. 6E, 6F**). However, no significant difference was observed in the proportion of GATA3^+^ CD11a^hi^CD44^+^ T cells, which were only weakly induced independent of genotype (**Fig. 6G**). Of note, we did not find any statistical difference in the absolute T cell numbers in the dLNs of d28 infected *Cd4^+/+^* and *Cd4^E4mΔ/Δ/ E4aΔ/Δ^* mice (Figure S6B). Since a transient IL-4 producing population has been observed early during infection with *L.major* in C57BL/6 mice [33], we examined effector T cells in the dLNs at 9 d.p.i. However, we did not observe a skewed GATA3^+^ CD11a^hi^CD44^+^ population in *Cd4^E4mΔ/Δ/ E4aΔ/Δ^* mice (**Fig. 6H, 6I**). Interestingly, upon *ex vivo* stimulation of CD11a^hi^CD44^+^ T cells from the dLNs of infected mice with soluble leishmania antigen (SLA), IFN-*γ* levels among CD8^-^CD11a^hi^CD44^+^ helper T cells from Cd4^+/+^ and *Cd4^E4mΔ/Δ/ E4aΔ/Δ^* mice were statistically similar (**Fig. 6J, Supplementary** **Fig.6C**). The same held true following restimulation with PMA/Ionomycin, (**Fig. 6K**). Local assessment of both IFN-*γ* (**Fig. 6L**) and IL-4 (**Supplementary Fig. 6D**) in the footpads of CD4^+/+^ and *Cd4^E4mΔ/Δ/ E4aΔ/Δ^* mice also revealed no significant differences. In contrast, increased levels of IL-1β, TNF and IL-10 in the footpads of *Cd4^E4mΔ/Δ/ E4aΔ/Δ^* infected mice were observed compared to controls (**Fig. 6M, 6N, 6O).** Taken together our studies highlights the importance of stable CD4 expression in supporting the differentiation of Th1 effector cells during *L.major* infection in C57BL/6 mice.

**Fig. 6:**
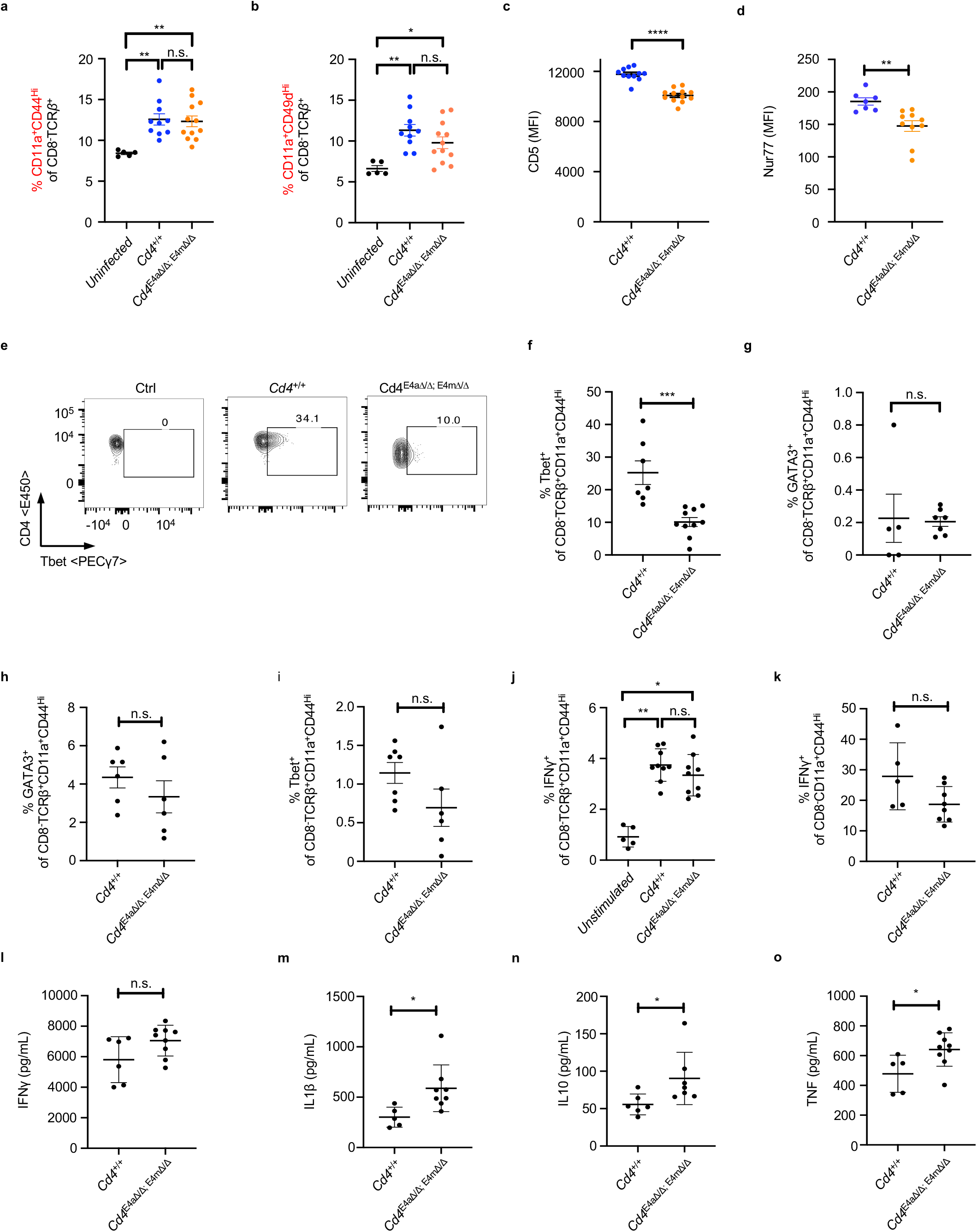
Reduced CD4 expression impairs the differentiation of Th1 cells during Leishmaniasis. **a,** Proportions of CD11a^+^CD44^hi^ T cells among CD8^-^ TCRβ^+^ T cells in the inguinal dLNs of mice, 9 days p.i. Data shown is mean ± SEM of 9-10 mice/group and is a summary of 2 independent experiments **p< 0.01 (One-way ANOVA and Kruskal Wallis test)**. b,** Proportions of CD11a^+^CD49d^hi^ T cells among CD8^-^ TCRβ^+^ T cells in the inguinal dLNs of mice, 9 days p.i. Data shown is mean ± SEM of 9-10 mice/group and is a summary of 2 independent experiments *p<0.05, **p< 0.01 (One-way ANOVA and Kruskal Wallis test)**. c,** CD5 MFI on CD11a^+^CD44^hi^ T cells among CD8^-^ TCRβ^+^ T cells in the inguinal dLNs of mice, 9 days p.i. Data shown is mean ± SEM of 9-10 mice/group and is a summary of 2 independent experiments ****p<p.0001 (Mann-Whitney test)**. d,** Nur77 MFI on CD11a^+^CD44^hi^ T cells among CD8^-^ TCRβ^+^ T cells in the inguinal dLNs of mice, 9 days p.i. Data shown is mean ± SEM of 7-10 mice/group and is a summary of 2 independent experiments ***p<0.001 (Mann-Whitney test)**. e,** FACS contour plot showing Tbet expression among CD4^+^CD8^-^ TCRβ^+^ T cells from the popliteal dLNs of mice, 28 days p.i. Data shown is representative of >3 independent experiments with 4-5 mice/group/experiment**. f,** Proportion of Tbet+ cells among CD8^-^ TCRβ^+^ CD11a^+^CD44^hi^T cells from the popliteal dLNs of mice, 28 days p.i. Data shown is a summary of 2 independent experiments with 4-5 mice/group/experiment. ***p<0.001 (Mann-Whitney test)**. g,** Proportion of GATA3+ cells among CD8^-^ TCRβ^+^ CD11a^+^CD44^hi^T cells from the popliteal dLNs of mice, 28 days p.i. Data shown is a summary of 2 independent experiments with 4-5 mice/group/experiment. (Mann-Whitney test)**. h,** Proportion of GATA3+ cells among CD8^-^ TCRβ^+^ CD11a^+^CD44^hi^T cells from the popliteal LNs of mice, 9 days p.i. Data shown is a representative of 2 independent experiments with 4-5 mice/group. (Mann-Whitney test)**. i,** Proportion of Tbet+ cells among CD8^-^ TCRβ^+^ CD11a^+^CD44^hi^T cells from the popliteal dLNs of mice, 9 days p.i. Data shown is a representative of 2 independent experiments with 6 mice/group. (Mann-Whitney test)**. j,** Proportion of IFN*γ*+ cells among CD8^-^ TCRβ^+^ CD11a^+^CD44^hi^T cells from the inguinal dLNs of d28 infected mice after restimulation overnight with SLA *in vitro*. Data shown is representative of 2 independent experiments with 4-5 mice/group with technical replicates. *p<0.05, **p<0.01 (One-way ANOVA and Kruskal-Wallis test)**. k,** Proportion of IFN*γ*+ cells among CD8^-^ TCRβ^+^ CD11a^+^CD44^hi^T cells from the inguinal dLNs of d28 *L.major-*infected mice after restimulation with PMA/Ionomycin for 4 hrs. Data shown is representative of 2 independent experiments with 4-5 mice/group *p<0.05, **p<0.01 (One-way ANOVA and Kruskal-Wallis test)**. l-o,** Supernatants from homogenized footpads of infected mice 28d.p.i were analyzed by multiplex ELISA for the presence of IFN*γ*+ (**l**), IL-1β (**m**), IL-10 (**n**), and TNF-α (**o**). (5-6 mice/group) *p<0.05 (Mann-Whitney test)

### TET-mediated demethylation during thymic development is critical for optimal gene function in effector T cells

To identify genes that share common regulatory features with *Cd4*, we next employed a genome-wide analysis using publicly available datasets for ATAC-Seq, H3K27Ac ChIP-Seq, RNA Seq and global 5hmC profiling by cytosine-5-methylenesulfonate immunoprecipitation (CMS-IP) to assemble a list of genes expressed in developing DP and CD4^+^ thymocytes with putative CREs that undergo DNA demethylation. We hypothesized that CREs active during thymic development would be critical in coordinating DNA demethylation and that lack of demethylation would impair gene expression in effector T cells by impairing the activity of stimulus-responsive CREs.

Of the 6719 genes that underwent a fold change >2 in gene expression from DN3 to CD4 single positive (SP) T cell specification in the thymus, 409 genes (10.8%) demonstrated active demethylation in CD4^+^ T cells, as assessed by presence of 5hmC detected via CMS-IP by Tsagaratou al.[6] (**Supplementary Fig. 7A**). 81.4 % of those genes (333 genes) showed a positive correlation with increased gene expression in CD4 SP cells and presence of 5hmC (**Supplementary Fig. 7A, Supplementary Table1**). Interestingly, a fraction of these genes (24% - 99 genes) displayed higher levels of 5hmC marks in DP precursors while most genes showed a further gain in 5hmC in CD4 SP T cells (**Supplementary Fig. 7B, Supplementary Table2**), suggesting that the demethylation process is initiated early during commitment to the CD4 lineage, congruent with our previous results [7]. Of the 350 genes that upregulated gene expression from DN3 to CD4 SP T cells (fold change > 2) and exhibited 5hmC in DP or CD4 SP thymic T cells, a striking 94 % (330 genes) had open chromatin peaks and concomitant H3K27Ac marks present in their gene bodies or upstream of their annotated TSS, highlighting an intimate link between CRE activity and presence of 5hmC (**Fig. 7A, Supplementary Table 3**).

**Fig. 7:**
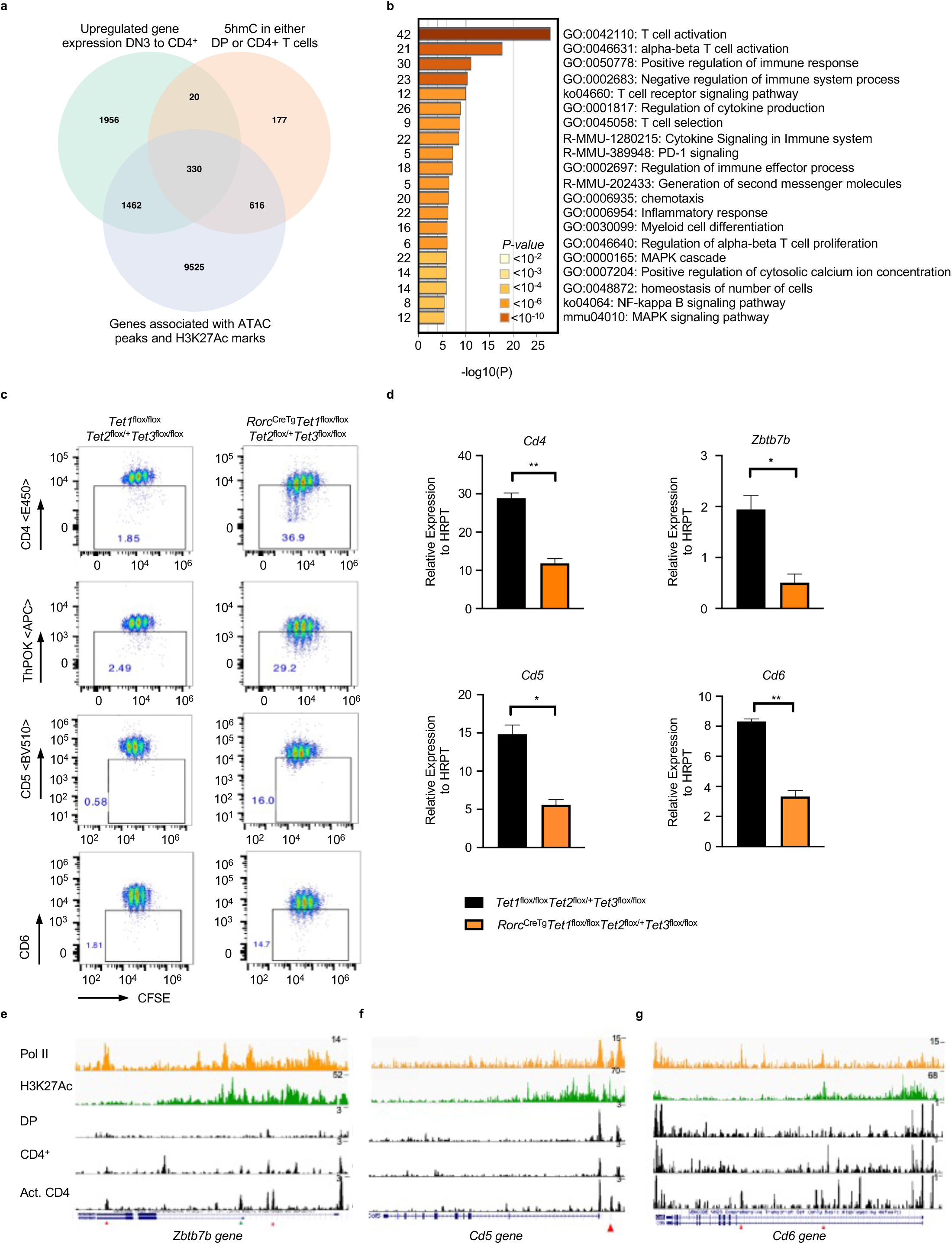
TET-mediated demethylation during thymic development is critical for optimal gene function in effector T cells. **a,** Venn diagram showing the number of genes that upregulate gene expression (a fold change >2 in gene expression from DN3 to CD4^+^SP) intersected with genes that have 5hmC in either DP or CD4^+^ SP T cells (Intragenic 5hmC log2 CMS-IP/Input >2) intersected with genes containing open chromatin ATAC-Seq peaks and H3K27Ac marks in either DP or CD4+ T cells. **b,** Metascape bar graph showing top non-redundant enrichment clusters among genes that have novel accessibility ATAC-Seq in activated T cells, undergo DNA demethylation and upregulate RNA expression during thymic development. Number of genes in each cluster is indicated on the left and cluster IDs are shown on the right. Bar graph was generated using the Metascape gene annotation and analysis online resource. List of genes used for the enrichment analysis is found in Supplementary Table 4. **c,** FACS plot showing CD4, ThPOK, CD5 and CD6 expression in CFSE-labeled T cells from control or RorcCre^Tg^ Tet1^fl/fl^ Tet2^fl/+^ Tet3^fl/fl^ mice 72 hrs post activation. Naïve CD4 T cells were FACS-sorted and activated with anti-CD3/CD28. Data is a representative of 2 experiments with at least 2 animals/genotype/experiment. **d,** *Cd4, Zbtb7b, Cd5 and Cd6* mRNA expression in control or RorcCre^Tg^ Tet1^fl/fl^ Tet2^fl/+^ Tet3^fl/fl^ T cells activated *in vitro* for 96 hrs with anti-CD3/CD28. Data shown is mean ± SEM of 2 mice/group and is representative of 2 independent experiments. * p<0.05, **p<0.01 (Student’s t test)**. e,** IGV snapshots of the *Zbtb7b* locus depicting p300 and H3K27Ac signals by ChIP-Seq in activated T (Th0) cells and ATAC Seq peaks in DP, CD4^+^ thymocytes and Th0 CD4^+^ T cells. Red arrows denote currently undefined CREs. Green arrow shows the location of a previously validated CRE PE in CD4^+^ T cells [58, 60]. **f,** IGV snapshot of the *Cd5* locus depicting p300 and H3K27Ac signals by ChIP-Seq in activated T (Th0) cells and ATAC Seq peaks in DP, CD4^+^ thymocytes and Th0 CD4^+^ T cells. Red arrows denote currently undefined CREs. **g,** IGV snapshot of the *Cd6* locus depicting p300 and H3K27Ac signals by ChIP-Seq in activated T (Th0) cells and ATAC Seq peaks in DP, CD4^+^ thymocytes and activated CD4^+^ T cells. Red arrows denote currently undefined CREs.

We next manually curated the list of 350 genes to identify genes that contain new accessible peaks when activated *in vitro* (Th0 cells), similarly to *Cd4*. Strikingly, 44% among those genes had novel accessible peaks downstream or upstream of their annotated TSS in activated CD4^+^ T cells (**Supplementary Fig. 7C, Supplementary Table 4**). Gene ontology and enrichment analysis revealed that a significant proportion of these genes had fundamental roles in regulating TCR activation, signaling and differentiation pathways (**Fig. 7B**). Indeed, many of these genes such as *Cd5*, *Cd6, Zbtb7b* have been shown to modulate critical post-thymic functions in helper T cells [34–39], suggesting that their epigenetic programming during development may be critical for their optimal expression in effector T cells. To assess the consequence of impaired thymic DNA demethylation, we generated *Rorc(t)^CreTg^ Tet1^fl/fl^ Tet2^fl/+^ Tet3^fl/fl^* mice in which excision of TET alleles occurs during the transition of immature single positive (ISP) CD8^+^ to DP thymocytes [40, 41]. We reasoned that deletion of one allele of TET2 may allow for recovery of genes that undergo demethylation in a TET2-dependent manner but without impairing homeostasis, as akin to *CD4^CreTg^ Tet 2/3* floxed mice, *Rorc(t)^CreTg^ Tet1/2/3* compound floxed mice have profound developmental defects, display severe autoimmune phenotypes with an onset of 4-5 weeks and enlarged spleens and lymph nodes [42] (**Supplementary Fig. 7D**), making the assessment of post-thymic effector T cells difficult. In contrast, *Rorc(t)^CreTg^ Tet1^fl/fl^ Tet2^fl/+^ Tet3^fl/fl^* mice were healthy, displayed no splenomegaly or lympho-adenopathies and had equivalent proportions of naïve CD44^-^CD62L^+^ T lymphocyte compartments as control mice (**Supplementary Fig. 7E**). Furthermore, naïve CD4 T cells from *Rorc(t)^CreTg^ Tet1^fl/fl^ Tet2^fl/+^ Tet3^fl/fl^* mice had comparable CD4 expression to controls (**Supplementary Fig. 7F**). However, upon T cell proliferation, CD4 expression was dramatically reduced (**Fig. 7C, Supplementary Fig. 7G**). In a similar pattern, ThPOK (*Zbtb7b* gene), CD5 and CD6 protein and mRNA expression were significantly reduced following proliferation (**Fig.7C, 7D, Supplementary** **Fig.7G**). Upon co-transfer of naïve *Rorc(t)^CreTg^ Tet1^fl/fl^ Tet2^fl/+^ Tet3^fl/fl^* and CD45.1 CD4 T cells into *Rag^-/-^* mice, there was significant downregulation of CD5, CD6 and ThPOK expression at 7 days post transfer (**Supplementary** **Fig.7H**). Of note, a reduction in CD5 and CD6 expression on naïve T cells was also seen prior to activation (**Supplementary Fig. 7F)**, suggesting a role for DNA demethylation in the regulation of these genes in the periphery. Nonetheless, these were significantly further downregulated following replication (**Fig. 7C,7D, Supplementary Fig. 7G**). *Cd5*, *Cd6* and *Zbtb7b* loci displayed novel chromatin accessible sites in activated CD4^+^ T cells (red arrows) which coincided with presence of H3K27Ac and/or presence of Pol II marks, suggestive of putative CREs. (**Fig. 7E, 7F, 7G**). Together our findings highlight the conservation of similar patterns of epigenetic regulation as *Cd4* in a cohort of genes and demonstrates the importance of DNA demethylation during development in ensuring proper gene function in effector helper T cells.

## DISCUSSION

The role of DNA methylation in intragenic and intergenic regions of the genome remain ill-defined. In this study, using the *Cd4* gene as a model locus, we demonstrated mechanistically how DNA methylation in non-promoter regions affects gene expression and illustrated the biological importance of DNA demethylation during thymic development.

DNA demethylation during T cell development sculpts the chromatin landscape for the subsequent function of stimulus-responsive regulatory elements in peripheral effector T cells. We showed that a novel stimulus-responsive CRE, E4a, acts in a partially redundant manner with a developmental CRE, E4m, to maintain CD4 expression in effector T cells. In the absence of TET1/3-mediated DNA demethylation in the thymus, the functions of E4a/E4m were compromised in effector cells and resulted in a loss of CD4 expression during cell proliferation. This loss strongly correlated with *de novo* DNA methylation at the *Cd4* promoter and gene expression could be rescued by knocking down *Dnmt1* and *Dnmt3a*. *De novo* methylation was a result of reduced transcriptional activity and reduced promoter H3K4me3 levels, as boosting enhancer function restored CD4 expression during replication and prevented gain of new methylation at the *Cd4* promoter by maintaining H3K4me3. Together these findings highlight a novel role for DNA demethylation in enabling the proper function of promoter/enhancer activities to maintain gene expression and H3K4me3 marks, and in preventing spurious *de novo* DNA methylation.

Our findings are in agreement with the established antagonistic relationship between H3K4me3 and *de novo* DNA methylation [21, 43, 44]. In a previous study using a GAL-4/ TATA-binding protein tethering system, it was elegantly shown that the positioning of the transcription machinery before implantation determines what regions will be protected from subsequent *de novo* methylation in blastocysts [45] . Transcription factors and the RNA polymerase complex were shown to play a major role in protecting recognized regions from *de novo* methylation by recruiting the H3K4 methylation machinery [45]. Our findings add to the earlier study, suggesting that the fidelity of maintenance of existing DNA methylation patterns in somatic cells and/or protection against rampant *de novo* methylation during replication are critically dependent on H3K4me3 signatures. It is enticing to speculate that such mechanisms may be involved in some of the unknown etiologies of promoter hypermethylation of a cohort of genes in certain disorders such as cancer [46, 47], although future work is needed.

While we did not find a role for E4a activity during thymic development, the intergenic space upstream of the *Cd4* locus in *Cd4^E4aΔ/Δ^* and *Cd4^E4mΔ/Δ/ E4aΔ/Δ^* T cells contained significant DNA methylation. We speculate that E4a may have a function in coordinating TET1/3-mediated demethylation together with E4m and E4p [8] or that it has a regulatory role in preventing spurious DNMT3a/3b activity in this region [3, 48–51]. In addition, we have documented a contribution for the CD4 co-receptor levels in modulating effector T cell differentiation. Beyond its critical role in regulating TCR signaling during thymic T cell commitment [8, 52], the importance of maintaining high CD4 expression during the differentiation of effector T cells remained unclear. We showed here that elevated CD4 expression was critical for optimal TCR signaling and promoted differentiation into Tbet-expressing Th1 lineage T cells in a mouse model of Leishmaniasis. Our data support the idea that strength of TCR signaling impacts effector T cell differentiation but does not skew their polarization into alternate subsets, a phenomenon most likely driven by environmental signals such as cytokines. Interestingly, we did not find a significant defect in IFN-*γ* production. It is noteworthy that due to the dramatic loss of CD4 during replication in *Cd4^E4mΔ/Δ/ E4aΔ/Δ^* mice, helper T cells had to be identified based on a TCRβ^+^ CD8a^-^ gating strategy. However, it has previously been reported that a population of MHC-Class II-restricted DN T cells (TCRβ^+^CD4^-^CD8^-^), which are high producers of IFN-*γ*, is induced during *L.major* infection [53, 54]. In a *Trichinella spiralis* model, these DN T cells have been shown to produce IFN-*γ* in a Tbet-independent manner [55]. Thus, we cannot exclude a role for CD4 in the skewing of the DN population in our model. In addition, previous studies have shown that other CD4^+^ T cell subsets, including regulatory T cells, play a role in *L.major* parasite persistence and clearance [56, 57] and therefore the role of CD4 in the differentiation of these subsets in this model cannot be excluded.

Lastly, building on principles from genomic studies of the *Cd4* locus, we have identified a cohort of genes whose expression in effector T cells was dependent on developmental programming. Notably, the *Zbtb7b* gene shares similar characteristics with *Cd4*. *Zbtb7b* expression is modulated during positive selection and differentiation of MHC-Class II-selected T cells in the thymus via two CREs, TE and PE [58–60]. Ablation of PE and TE abolishes *Zbtb7b* expression during thymic differentiation [61]. It is not known if PE and TE coordinate the demethylation process akin to the *Cd4* CRE*s*. Another CRE, general T cell element (GTE) situated between PE and TE has been reported to have enhancer properties in CD4^+^ T cells, although its endogenous role has not been tested [59]. Here, we have found enhanced chromatin accessibility in a putative CRE proximal to the GTE region in effector CD4^+^ T cells. Given the proliferation-associated loss of expression of *Zbtb7b* in T cells deprived of Tet activity during development, we speculate that GTE may be a stimulus-responsive element sensitive to intragenic or intergenic DNA methylation. Notably, in this study our analysis is confined to integration of genes with accessibility peaks early during early T cell activation (Th0 conditions), whereas many genes may require additional differentiation signals for optimal expression. Future experiments are warranted to delineate CREs that modulate expression in differentiated effector subsets.

These studies will be instrumental as roughly 90% of autoimmunity-associated genetic variants identified via genome-wide association studies lie in noncoding regions of the genome. It was recently reported that stimulation-responsive chromatin regions correlate with significant trait heritability in multiple immune cell types, pointing to potential causality of stimulus-responsive CREs in autoimmune diseases [62, 63]. Thus, viewed in this perspective, our study invokes a putative link between developmental epigenetic processes in shaping the activity of these stimulus-responsive elements in effector T cells.

## METHODS

### Mice

F0 mice were genotyped by PCR amplication of the targeted region followed by TA cloning, and Sanger sequencing of individual colonies. Founders bearing E4a deletions in chromosome 6 with MM10 coordinates 124901815 – 124902199 were then backcrossed at least two times to wild-type C57BL/6J mice. For the generation of *Cd4*^E4mΔ/Δ E4aΔ/Δ^ mutant mice, *Cd4*^E4aΔ/Δ^ embryos were used for CRISPR injections using the same guide RNAs used for the generation of *Cd4*^E4aΔ/Δ^ mutants. The generation of *Cd4*^E4mΔ/Δ^ mice has been described previously[8]. *Cd4*^E4mflox/flox^ mice were also generated via CRISPR-Cas9. Briefly, two sgRNAs targeting the E4m region and donor single-stranded DNA oligos containing loxP sites with 60bp homology to sequences on each side of the sgRNA target sites were injected into C57BL/6J zygotes. To facilitate detection of correct insertions, the donor template oligos were engineered to contain an Nhe1 restriction site on the 5’ arm and an EcoR1 site on the 3’ arm. Integration of loxP sites were first assessed by PCR amplification of the expected modified region followed by enzymatic digestion of PCR products by EcoR1 or NHE1. After determination of loxP sites on the same allele of E4m, respective homologous regions bearing the loxP sites were PCR amplified, TA cloned and Sanger sequenced to ensure no mutations around integrated loxP sites.

*Tet1* and *Tet2* floxed (C57BL/6) and *Tet3* floxed mice (129 backcrossed to C57BL/6) were kindly provided by Dr. Iannis Aifantis and Dr. Yi Zhang, respectively. These mice were then backcrossed onto *RorcCre*^Tg^ (B6.FVB-Tg(Rorc-cre)1Litt/J #022791) mice. B6.SJL-CD45.1 mice were purchased from the Jackson Laboratory (#002014). B6.129S7-Rag1tm1Mom/J were purchased from the Jackson Laboratory (#002216). All mice were maintained under specific pathogen-free conditions at the barrier animal facility at University of Iowa Carver College of Medicine. All experiments were performed in accordance with the protocol approved by the IACUC at the University of Iowa Carver College of Medicine.

### FACS sorting

T cells were enriched from the lymph nodes and spleen of mice using the Dynabeads untouched mouse T cell kit (Thermo Fisher Scientific) or the CD4+ T cell selection kit (Miltenyi). Naïve CD4+ T cells were then isolated by flow cytometry based on the markers CD4+CD62L+CD44−CD25− on a FACS Aria II or FACS Fusion Sorter.

### Antibodies and Flow Cytometry

The following antibodies from Tonbo, Biolegend or ThermoFisher Scientific were used: CD62L (MEL-14), CD25 (PC61.5), CD44 (IM7), CD19 (ID3), CD5 (53-7.3), CD6 (OX-129), Zbtb7b (T43-94), CD4 (RM4-5), CD8a (53-6.7), Nur77 (12.14), CD11a (M17/4), CD49d (R1-2), TNF (MP6-XT22), IFN-*γ* (XMG1.2), CD45.2 (104), CD45.1 (A20), GATA3 (TWAJ), Tbet (4B10). Ghost Dye from Tonbo was used to gate out dead cells from all analysis. LN and spleen were isolated by dissection from mice and then mashed through a 70-μm filter. Red Blood Cells were lysed in ACK lysis buffer (Lonza), counted and stained in 2% IMDM (Gibco) with a Live/Dead exclusion Ghost Dye (Tonbo), followed by staining with surface markers for 30 min at 4 degrees. Details of all antibodies are provided in the supplementary method section. For transcription factor staining, cells were fixed overnight in the Foxp3/Transcription Factor/Fixation-Concentrate kit (Tonbo) after surface staining. After fixation, cells were permeabilized and stained with the appropriate antibodies in PBS for 1hr at room temperature. For cytokine analysis, after surface staining cells were fixed at a final concentration of 2% PFA for 15 min at room temperature, followed by 2 washes with PBS and overnight storage. The next day, cells were permeabilized and stained for 1hr at room temp for cytokines. Stained cells were analyzed on a Cytoflex flow analyzer or an LSR II flow analyzer.

### T cell activation and retroviral transduction

Tissue culture plates were coated with polyclonal goat anti-hamster IgG (MP Biomedical) at 37 °C for at least 2 h or overnight at 4°C and washed 2× with PBS before cell plating. FACS-sorted CD4^+^CD8^−^CD25^−^CD62L^+^CD44^lo^ naïve T cells were seeded in T cell medium [RPMI 1640 (Gibco), 10% heat-inactivated FBS (Atlantis), 2 mM L-glutamine, 50 µg/ml gentamicin, 1% Penn/Strep, 50uM 2-mercaptoethanol (Gibco)] along with anti-CD3 (BioXcell, clone 145-2C11, 0.25 μg/ml) and anti-CD28 (BioXcell, clone 37.5.1, 1 μg/ml) antibodies. Forty-eight hours later cells were lifted off the plates and cultures were supplemented with 100 U/ml recombinant human IL-2 (Peprotech). CFSE labeling for certain experiments was performed prior to seeding on tissue culture plates.

For *Dnmt1* and *Dnmt3a* knockdowns, shRNA plasmids on a mIR-E backbone were provided by Johannes Zuber [64]; MSCV-beta-catenin-IRES-GFP was a gift from Tannishtha Reya (Addgene plasmid #14717). MSCV-Cre-IRES-GFP was previously described [9]. Retroviruses were packaged in PlatE cells by transient transfection using TransIT 293 (Mirus Bio) or Lipofectamine 3000 (Thermo Fisher). Cells were transduced by spin infection at 1200 × *g* at 32 °C for 90 min in the presence of 10 μg/ml polybrene (Santa Cruz) 12–16 h post-activation with anti-CD3/anti-CD28. Viral supernatants were removed the next day and replaced with fresh medium containing anti-CD3/anti-CD28. Cells were lifted off 24 h later and supplemented with 100 U/ml recombinant human IL-2.

### Methylation analysis

Genomic DNA was isolated from FACS-sorted T cell populations using genomic DNA isolation kits (Qiagen). Purified genomic DNA was subjected to bisulfite treatment with the Qiagen EpiTect bisulfite kit. For locus-wide bisulfite sequencing, CATCH-seq was performed as previously described[7, 13] using BAC clone RP24-330J12 (BACPAC Resource Center, CHORI).

### Low input histone ChIP-qPCR

100,000-150,000 T cells were isolated by cell sorting and processed for ultralow input micrococcal nuclease-based native ChIP (ULI-NChIP) [65]. For immunoprecipitation, 2 µg of antibody was used per reaction. H3K9me3 (Lot. A2217P) and H3K4me3 (Lot No. A1052D) antibodies were from Diagenode. Immunoprecipitated and input DNA were extracted by phenol-chloroform extraction and used for qPCR with SsoAdvanced Universal SYBR Green supermix (Biorad). All primer sequences used are listed in **Supplementary Table 5**.

### Real-time quantitative PCR

DNase I-treated total RNA was prepared from sorted T cells using RNeasy RNA isolation kit (Qiagen) and cDNA was synthesized using an iScript cDNA synthesis kit (Biorad). Quantitative PCR was performed using SsoAdvanced Universal SYBR Green supermix (Biorad) and a CFX Connect Real-time PCR detection system (Biorad). List of primers used are detailed in the supplementary table. All primer sequences used are listed in **Supplementary Table 5.**

### Rag Knockout T cell transfer

Sex and age-matched CD4+ T cells were enriched from CD45.1 and naïve control or RorcCre Tet1/3 mice using the Dynabeads untouched CD4 T cell kit (ThermoFisher Scientific), ensuring >85% purity. Cells were counted and CFSE labeled and mixed at a 1:1 ratio before transfer by i.v injections into 6-8 weeks old *Rag-/-* recipients. In some experiments, naïve CD4 T cells were FACS-sorted after enrichment with Dynabeads.

### Leishmania infection

The *L. major* strain WHOM/IR/-173 was grown *in vitro* in complete M199 media supplemented with 5% HEPES, 10% FBS, and 1% penicillin-streptomycin at 26°C. Metacyclic promastigotes were isolated from stationary phase cultures and enriched using a Ficoll gradient as described previously [66]. Mice were infected with 2×10^6^ *L. major* promastigotes per footpad in a volume of 50 μl PBS. Footpad measurements were taken weekly and prior to infection by caliper measurement. For quantification of *L. major* promastigotes, footpads were surgically dissected and homogenized by douncing as previously described [66], and 10-fold serial limiting dilutions to extinction of the homogenates were plated in triplicates in 96-well flat-bottomed plates in complete M199 media. Four to 6 days after culture at 26°C, each well was analyzed under a microscope for the presence or absence of *L. major* to determine the titers. Media only controls were included to ensure no cross contamination. Promastigotes were counted in a blinded manner at the lowest dilution factor and triplicates were averaged.

### Multiplex cytokine analysis

Homogenates collected after douncing of tissue from the footpads of infected animals were spun down to remove cellular components and supernatants were collected and stored at - 80°C for cytokine measurements using a Th1-Th2 mouse ProcartaPlex Panel (Thermo Fisher Scientific). 50ul of supernatants with appropriate dilutions were used for cytokine estimations and fluorescent readings were acquired on a BioRad Bio-Plex (Luminex 200) instrument.

### *In vitro* restimulation and flow cytometry

Draining popliteal or inguinal lymph nodes were harvested from infected and uninfected mice, and single-cell suspensions were made by mechanical dissociation. Cells were then seeded at 2×10^5^ cells/well in 200 µl of complete RPMI medium [RPMI 1640, 10% heat-inactivated FBS, 2 mM L-glutamine, 50 µg/ml gentamicin, 50uM 2-mercaptoethanol], in round bottomed 96-well plates with or without stimuli. Cells were then stimulated with PMA/Ionomycin (Tonbo) for 4 hours in the presence of Monensin and Brefeldin A (Tonbo) followed by staining for flow cytometry. For restimulation with soluble Leishmania antigen (SLA), lymph nodes were enzymatically digested with 100U/mL of collagenase type II (Gibco) at 37°C for 30 min to liberate lymphocytes and antigen-presenting cell populations, after which cells were dispersed through a 100 micron nylon mesh. SLA was prepared by repeated freeze-thaw cycles of late-log phase promastigotes of *L.major* grown in liquid culture and protein concentrations were estimated by the Bradford method as described previously [67]. 10 µg/ml total SLA lysate or PBS was added to plated cells for 18 hrs and Monensin/Brefeldin A was then added 6hrs before processing for intracellular cytokines measurements.

### Bioinformatics processing and analysis

We used our computational pipeline developed with Nextflow (https://www.nextflow.io) to integrate gene expression data with ChIP-Seq and ATAC-Seq data in order to generate a prioritized list of target genes. The change in gene expression (DN3 to CD4+) was assessed using publicly available RNA sequencing data from NCBI-GEO. The processed gene counts table for dataset GSE109125 was imported into R and the raw counts were normalized using the DESeq2 package. Log_2_ fold changes in gene expression were computed using the Wald test in DESeq2, and genes with counts less than 10 in all samples were removed from further analysis. The results of the differential expression analysis were visualized in R with customized volcano plots generated using the ggplot2 graphic package. Processed data from CMS-IP experiments performed in thymocytes were downloaded from Tsagaratou et al [6] . To identify genomic locations with acetylated histone marks (H3K27), we analyzed publicly available ChIP-Seq data from the NCBI SRA archive. The datasets (SRR5385354, SRR5385355, SRR5385352, and SRR5385353) were downloaded using the SRA toolkit and the SRA files were converted to fastq files for processing. The raw sequences were first trimmed with fastp and then aligned to the mouse genome (version mm10) using bwa mem. Peak calling was performed with Genrich using the following filtering options: removal of PCR duplicates and retention of unpaired alignments. The default p-value of 0.01 was used for statistical significance. Processed ATAC-Seq data containing peak location information for immune cells was downloaded from the Immgen project. Peak files from both the ChIP-Seq and ATAC-Seq analyses were imported into R and analyzed in a similar way using the ChIPseeker package. Association of peaks with genes was done using the ChIPseeker annotatePeak function using gene location data from the UCSC known gene database. All of the filtering and integration of gene lists from the above 3 analyses were performed in R. The final list of genes with new accessible ATAC-Seq peaks in Th0 cells was then manually curated to ascertain presence of peaks within the gene-body and <10kb away from the nearest annotated gene TSS. Gene enrichment analysis was performed using the Metascape online platform[68]. p300 and H3K27Ac ChIP-Seq datasets in Th0 cells are from previously published work from Ciofani et al. (GSE40918)

## QUANTIFICATION AND STATISTICAL ANALYSIS

GraphPad Prism 8.0 software was used for data analysis. Data are represented as the mean ± SEM unless otherwise specified. Statistical significance was determined by the Mann-Whitney *U* test for comparison of 2 groups for *in vivo* analysis or by ANOVA, Student’s t-test or multiple comparison t tests for *in vitro* experiments as described in the Figure legends. A *p* value of less than 0.05 was considered statistically significant.

## DATA AVAILABILITY STATEMENT

Bisulfite datasets using CATCH-Seq will be deposited in the Sequence Read Archive (SRA) and accession codes will be provided in this manuscript before publication. All other data are available from the corresponding author upon reasonable request.

## ACKNOWLEDGEMENTS

Some of the data presented herein were obtained at the Flow Cytometry Facility, which is a Carver College of Medicine / Holden Comprehensive Cancer Center core research facility at the University of Iowa. The facility is funded through user fees and the generous financial support of the Carver College of Medicine, Holden Comprehensive Cancer Center, and Iowa City Veteran’s Administration Medical Center. We are grateful to Dr. Mary Wilson at the University of Iowa for insights on Leishmaniasis-related experiments, Dr. William Nauseef at the University of Iowa for insights on the manuscript, and Dr. Y. Zhang and Dr. I. Aifantis for sharing of TET mutant mice. We thank Dr. Sangyong Kim and the Rodent Genetic Engineering Core (RGEC) of NYU Medical Center for generation of the *Cd4* mutant mouse strains. The RGEC is partially supported by Cancer Center Support grant P30CA016087 at the Laura and Isaac Perlmutter Cancer Center of the NYU Medical Center. D.R.L. was supported by the Howard Hughes Medical Institute.

## AUTHOR CONTRIBUTIONS

AT, PP, KM designed and performed experiments, analyzed data and provided input to the manuscript. CA provided technical assistance in the creation and analysis of CRISPR founder mice at NYU. KD did methylation captures and bioinformatic analyses of *Cd4* locus-wide methylation captures; HK performed the ChIP-Seq alignments, bioinformatics analysis and integrated datasets for genome-wide analysis. RY provided initial insights on ATAC-Seq analysis. MY performed Leishmaniasis experiments with supervision and insights from PG. PG assisted in the interpretation of data and provided input to the manuscript. TM provided experimental support for multiplex assays and input to the manuscript. DRL provided guidance and support during the initial conception of this study. PDI designed, performed and supervised experiments throughout the study, co-analyzed data with AT, PP, KM and wrote the manuscript with input from DRL and co-authors.

## REFERENCES

1. Li, E. and Y. Zhang, DNA methylation in mammals. Cold Spring Harb Perspect Biol, 2014. 6(5): p. a019133.

2. Jones, P.A., Functions of DNA methylation: islands, start sites, gene bodies and beyond. Nat Rev Genet, 2012. 13(7): p. 484–92.

3. Baubec, T., et al., Genomic profiling of DNA methyltransferases reveals a role for DNMT3B in genic methylation. Nature, 2015. 520(7546): p. 243–7.

4. Wu, H., et al., Dnmt3a-dependent nonpromoter DNA methylation facilitates transcription of neurogenic genes. Science, 2010. 329(5990): p. 444–8.

5. Lister, R., et al., Human DNA methylomes at base resolution show widespread epigenomic differences. Nature, 2009. 462(7271): p. 315–22.

6. Tsagaratou, A., et al., Dissecting the dynamic changes of 5-hydroxymethylcytosine in T-cell development and differentiation. Proc Natl Acad Sci U S A, 2014. 111(32): p. E3306–15.

7. Sellars, M., et al., Regulation of DNA methylation dictates Cd4 expression during the development of helper and cytotoxic T cell lineages. Nat Immunol, 2015. 16(7): p. 746–54.

8. Issuree, P.D., et al., Stage-specific epigenetic regulation of CD4 expression by coordinated enhancer elements during T cell development. Nat Commun, 2018. 9(1): p. 3594.

9. Chong, M.M., et al., Epigenetic propagation of CD4 expression is established by the Cd4 proximal enhancer in helper T cells. Genes Dev, 2010. 24(7): p. 659–69.

10. Henson, D.M., et al., A silencer-proximal intronic region is required for sustained CD4 expression in postselection thymocytes. J Immunol, 2014. 192(10): p. 4620–7.

11. Kojo, S., et al., Runx-dependent and silencer-independent repression of a maturation enhancer in the Cd4 gene. Nat Commun, 2018. 9(1): p. 3593.

12. Creyghton, M.P., et al., Histone H3K27ac separates active from poised enhancers and predicts developmental state. Proc Natl Acad Sci U S A, 2010. 107(50): p. 21931–6.

13. Day, K., J. Song, and D. Absher, Targeted sequencing of large genomic regions with CATCH-Seq. PLoS One, 2014. 9(10): p. e111756.

14. Visel, A., et al., ChIP-seq accurately predicts tissue-specific activity of enhancers. Nature, 2009. 457(7231): p. 854–8.

15. Weinert, B.T., et al., Time-Resolved Analysis Reveals Rapid Dynamics and Broad Scope of the CBP/p300 Acetylome. Cell, 2018. 174(1): p. 231–244 e12.

16. Raisner, R., et al., Enhancer Activity Requires CBP/P300 Bromodomain-Dependent Histone H3K27 Acetylation. Cell Rep, 2018. 24(7): p. 1722–1729.

17. Sawada, S., et al., A lineage-specific transcriptional silencer regulates CD4 gene expression during T lymphocyte development. Cell, 1994. 77(6): p. 917–29.

18. Taniuchi, I., et al., Evidence for distinct CD4 silencer functions at different stages of thymocyte differentiation. Mol Cell, 2002. 10(5): p. 1083–96.

19. Zou, Y.R., et al., Epigenetic silencing of CD4 in T cells committed to the cytotoxic lineage. Nat Genet, 2001. 29(3): p. 332–6.

20. Guo, X., et al., Structural insight into autoinhibition and histone H3-induced activation of DNMT3A. Nature, 2015. 517(7536): p. 640–4.

21. Ooi, S.K., et al., DNMT3L connects unmethylated lysine 4 of histone H3 to de novo methylation of DNA. Nature, 2007. 448(7154): p. 714–7.

22. Itoh, Y., et al., Decreased CD4 expression by polarized T helper 2 cells contributes to suboptimal TCR-induced phosphorylation and reduced Ca2+ signaling. Eur J Immunol, 2005. 35(11): p. 3187–95.

23. Tubo, N.J. and M.K. Jenkins, TCR signal quantity and quality in CD4(+) T cell differentiation. Trends Immunol, 2014. 35(12): p. 591–596.

24. Bhattacharyya, N.D. and C.G. Feng, Regulation of T Helper Cell Fate by TCR Signal Strength. Front Immunol, 2020. 11: p. 624.

25. Sacks, D. and N. Noben-Trauth, The immunology of susceptibility and resistance to Leishmania major in mice. Nat Rev Immunol, 2002. 2(11): p. 845–58.

26. Christiaansen, A.F., et al., CD11a and CD49d enhance the detection of antigen-specific T cells following human vaccination. Vaccine, 2017. 35(33): p. 4255–4261.

27. McDermott, D.S. and S.M. Varga, Quantifying antigen-specific CD4 T cells during a viral infection: CD4 T cell responses are larger than we think. J Immunol, 2011. 187(11): p. 5568–76.

28. Hornick, E.E., Z.R. Zacharias, and K.L. Legge, Kinetics and Phenotype of the CD4 T Cell Response to Influenza Virus Infections. Front Immunol, 2019. 10: p. 2351.

29. Martin, D.L. and R.L. Tarleton, Antigen-specific T cells maintain an effector memory phenotype during persistent Trypanosoma cruzi infection. J Immunol, 2005. 174(3): p. 1594–601.

30. Rai, D., et al., Tracking the total CD8 T cell response to infection reveals substantial discordance in magnitude and kinetics between inbred and outbred hosts. J Immunol, 2009. 183(12): p. 7672–81.

31. Moran, A.E., et al., T cell receptor signal strength in Treg and iNKT cell development demonstrated by a novel fluorescent reporter mouse. J Exp Med, 2011. 208(6): p. 1279–89.

32. Azzam, H.S., et al., CD5 expression is developmentally regulated by T cell receptor (TCR) signals and TCR avidity. J Exp Med, 1998. 188(12): p. 2301–11.

33. Uzonna, J.E., K.L. Joyce, and P. Scott, Low dose Leishmania major promotes a transient T helper cell type 2 response that is down-regulated by interferon gamma-producing CD8+ T cells. J Exp Med, 2004. 199(11): p. 1559–66.

34. Vacchio, M.S., et al., A ThPOK-LRF transcriptional node maintains the integrity and effector potential of post-thymic CD4+ T cells. Nat Immunol, 2014. 15(10): p. 947–56.

35. Ciucci, T., et al., The Emergence and Functional Fitness of Memory CD4(+) T Cells Require the Transcription Factor Thpok. Immunity, 2019. 50(1): p. 91–105 e4.

36. Zinzow-Kramer, W.M., A. Weiss, and B.B. Au-Yeung, Adaptation by naive CD4(+) T cells to self-antigen-dependent TCR signaling induces functional heterogeneity and tolerance. Proc Natl Acad Sci U S A, 2019. 116(30): p. 15160–15169.

37. Henderson, J.G., et al., CD5 instructs extrathymic regulatory T cell development in response to self and tolerizing antigens. Immunity, 2015. 42(3): p. 471–83.

38. Kofler, D.M., et al., The Link Between CD6 and Autoimmunity: Genetic and Cellular Associations. Curr Drug Targets, 2016. 17(6): p. 651–65.

39. Kofler, D.M., et al., The CD6 multiple sclerosis susceptibility allele is associated with alterations in CD4+ T cell proliferation. J Immunol, 2011. 187(6): p. 3286–91.

40. Eberl, G. and D.R. Littman, Thymic origin of intestinal alphabeta T cells revealed by fate mapping of RORgammat+ cells. Science, 2004. 305(5681): p. 248–51.

41. Sun, Z., et al., Requirement for RORgamma in thymocyte survival and lymphoid organ development. Science, 2000. 288(5475): p. 2369–73.

42. Tsagaratou, A., et al., TET proteins regulate the lineage specification and TCR-mediated expansion of iNKT cells. Nat Immunol, 2017. 18(1): p. 45–53.

43. Cedar, H. and Y. Bergman, Linking DNA methylation and histone modification: patterns and paradigms. Nat Rev Genet, 2009. 10(5): p. 295–304.

44. Otani, J., et al., Structural basis for recognition of H3K4 methylation status by the DNA methyltransferase 3A ATRX-DNMT3-DNMT3L domain. EMBO Rep, 2009. 10(11): p. 1235–41.

45. Greenfield, R., et al., Role of transcription complexes in the formation of the basal methylation pattern in early development. Proc Natl Acad Sci U S A, 2018. 115(41): p. 10387–10391.

46. Herman, J.G. and S.B. Baylin, Gene silencing in cancer in association with promoter hypermethylation. N Engl J Med, 2003. 349(21): p. 2042–54.

47. Keshet, I., et al., Evidence for an instructive mechanism of de novo methylation in cancer cells. Nat Genet, 2006. 38(2): p. 149–53.

48. Jeong, M., et al., Large conserved domains of low DNA methylation maintained by Dnmt3a. Nat Genet, 2014. 46(1): p. 17–23.

49. Rinaldi, L., et al., Dnmt3a and Dnmt3b Associate with Enhancers to Regulate Human Epidermal Stem Cell Homeostasis. Cell Stem Cell, 2016. 19(4): p. 491–501.

50. Neri, F., et al., Intragenic DNA methylation prevents spurious transcription initiation. Nature, 2017. 543(7643): p. 72–77.

51. Weinberg, D.N., et al., The histone mark H3K36me2 recruits DNMT3A and shapes the intergenic DNA methylation landscape. Nature, 2019. 573(7773): p. 281–286.

52. Issuree, P.D., C.P. Ng, and D.R. Littman, Heritable Gene Regulation in the CD4:CD8 T Cell Lineage Choice. Front Immunol, 2017. 8: p. 291.

53. Mou, Z., et al., MHC class II restricted innate-like double negative T cells contribute to optimal primary and secondary immunity to Leishmania major. PLoS Pathog, 2014. 10(9): p. e1004396.

54. Antonelli, L.R., et al., Disparate immunoregulatory potentials for double-negative (CD4-CD8-) alpha beta and gamma delta T cells from human patients with cutaneous leishmaniasis. Infect Immun, 2006. 74(11): p. 6317–23.

55. Kannan, A.K., et al., T-Bet independent development of IFNgamma secreting natural T helper 1 cell population in the absence of Itk. Sci Rep, 2017. 7: p. 45935.

56. Anderson, C.F., S. Mendez, and D.L. Sacks, Nonhealing infection despite Th1 polarization produced by a strain of Leishmania major in C57BL/6 mice. J Immunol, 2005. 174(5): p. 2934–41.

57. Belkaid, Y., et al., CD4+CD25+ regulatory T cells control Leishmania major persistence and immunity. Nature, 2002. 420(6915): p. 502–7.

58. Muroi, S., et al., Cutting edge: fine-tuning of Thpok gene activation by an enhancer in close proximity to its own silencer. J Immunol, 2013. 190(4): p. 1397–401.

59. He, X., et al., CD4-CD8 lineage commitment is regulated by a silencer element at the ThPOK transcription-factor locus. Immunity, 2008. 28(3): p. 346–58.

60. Muroi, S., et al., Cascading suppression of transcriptional silencers by ThPOK seals helper T cell fate. Nat Immunol, 2008. 9(10): p. 1113–21.

61. Taniuchi, I., CD4 Helper and CD8 Cytotoxic T Cell Differentiation. Annu Rev Immunol, 2018. 36: p. 579–601.

62. Farh, K.K., et al., Genetic and epigenetic fine mapping of causal autoimmune disease variants. Nature, 2015. 518(7539): p. 337–43.

63. Calderon, D., et al., Landscape of stimulation-responsive chromatin across diverse human immune cells. Nat Genet, 2019. 51(10): p. 1494–1505.

64. Fellmann, C., et al., An optimized microRNA backbone for effective single-copy RNAi. Cell Rep, 2013. 5(6): p. 1704–13.

65. Brind’Amour, J., et al., An ultra-low-input native ChIP-seq protocol for genome-wide profiling of rare cell populations. Nat Commun, 2015. 6: p. 6033.

66. Sacks, D.L. and P.C. Melby, Animal models for the analysis of immune responses to leishmaniasis. Curr Protoc Immunol, 2015. 108: p. 19 2 1–19 2 24.

67. Coelho, E.A., et al., Immune responses induced by the Leishmania (Leishmania) donovani A2 antigen, but not by the LACK antigen, are protective against experimental Leishmania (Leishmania) amazonensis infection. Infect Immun, 2003. 71(7): p. 3988–94.

68. Zhou, Y., et al., Metascape provides a biologist-oriented resource for the analysis of systems-level datasets. Nat Commun, 2019. 10(1): p. 1523.

