## Supplementary Figures for "Epigenetic Programming during thymic development sets the stage for optimal function in effector T cells via DNA demethylation"

### SUPPLEMENTARY FIGURE LEGENDS

**Supplementary Fig.1: E4a modulates *Cd4* expression in effector T cells in a partially redundant manner with E4m**. **a**, DNA electrophoresis gel showing excision of the E4m allele after Cre-recombinase expression. Transduced T cells ($GFP^+$) were FACS-sorted 96hrs post-transduction and genomic DNA was isolated for PCR analysis. DNA from $Cd4^{E4m\Delta/\Delta}$ was used as a positive control. **b**, Dot Plot showing CD4 expression and CFSE dilution on control and $Cd4^{E4a\Delta/\Delta}$ CD4 T cells, 72hrs post activation with anti-CD3 and CD28. Experiment is representative of >3 experiments. **c**, Bar graph showing CD4 MFI on *in vitro* activated T cells from control or $Cd4^{E4a\Delta/\Delta}$ mice analyzed 72 hrs post activation. Cells were gated at equivalent cell division cycles. Data shown is representative > 3 experiments and expressed as mean ± SEM of 2 animals/group and 3 technical replicates/animal. ** p<0.01 (Student's t test). **d**, Bar graph showing CD4 gMFI on *in vitro* activated T cells from indicated genotypes analyzed 16 hrs and 48 hrs post activation. Data shown is a summary of 2 experiments and expressed as mean ± SEM of 2 animals/group/experiment. *p<0.05, ****p<0.0001 (2-Way ANOVA and Bonferroni test). **e**, CD4 mRNA expression (exon 1-2) in *in vitro* activated CD4 T cells with indicated genotypes. RNA was isolated 96hrs post activation. Data is expressed as mean ± SEM of 4-5 individual mice/group and is representative of two independent experiments. (One-Way ANOVA and Bonferroni test). **f**, CD4 mRNA expression (exon 2-3) in *in vitro* activated CD4 T cells with indicated genotypes. RNA was isolated 96hrs post activation. Data is expressed as mean ± SEM of 4-5 individual mice/group and is representative of two independent experiments. **g**, FACS dot plots showing CD4 expression and CFSE dilution at 72hrs on CD4 T cells following activation with indicated doses of anti-CD3 and 1ug/mL anti-CD28.  Data is representative of two independent experiments. **h**, CD4 gMFI expression on CD4 T cells 72hrs post activation with indicated concentrations of anti-CD3 and 1ug/mL anti-CD28. Cells were gated on equivalent cell division numbers. Data is expressed as mean ± SEM of 2 mice/group and is representative of two independent experiments. ***p<0.001 (2-way ANOVA with Tukey's multiple comparison test)

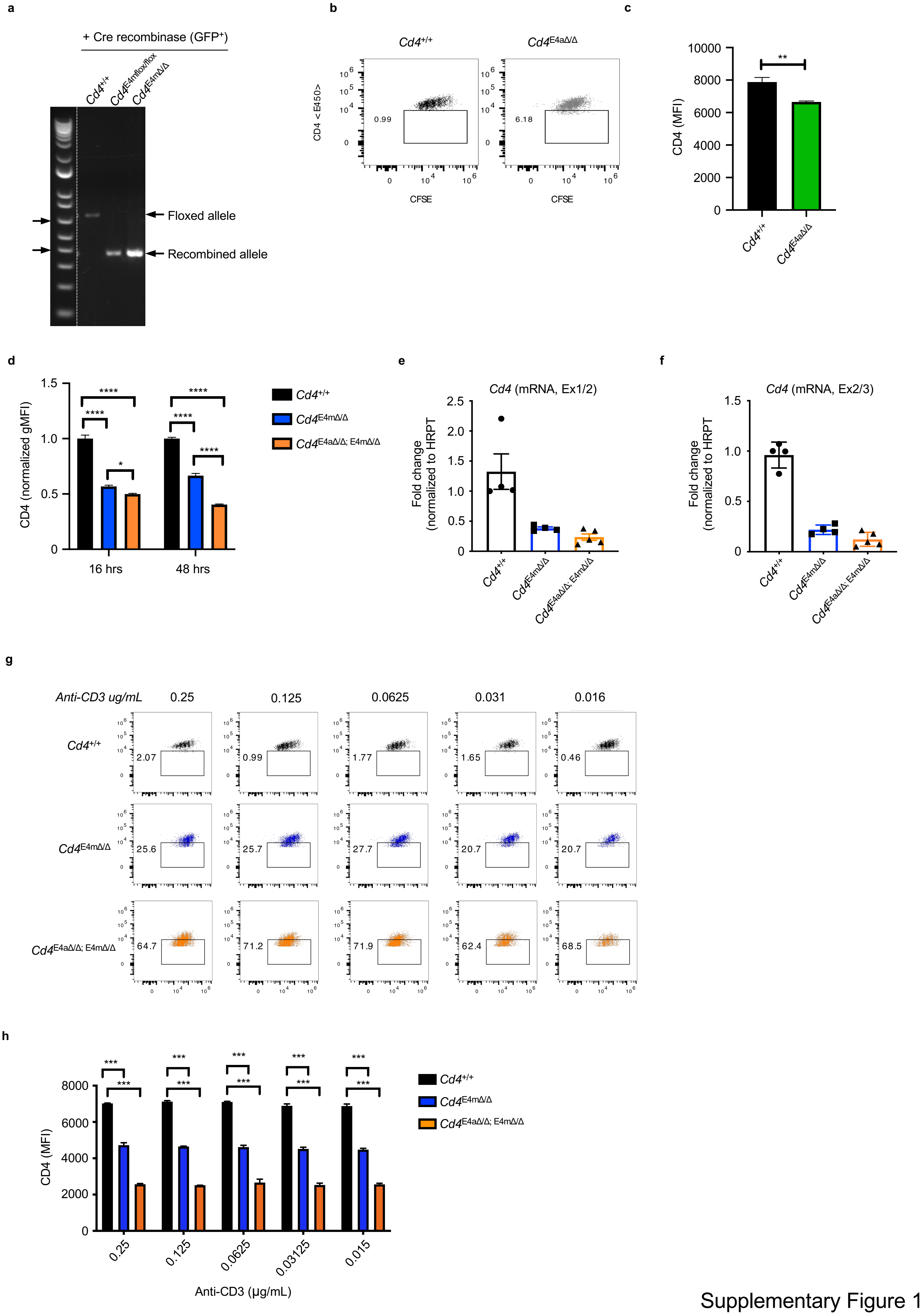


Supplementary Figure 1

**Supplementary Fig. 2: E4a is a stimulus-responsive *cis*-regulatory element licensed during development**. **a**, FACS contour data plots showing pre-selected TCR$\beta^{Lo}$CD69$^{-}$CD24$^{hi}$DP (top panel), recently selected TCR$\beta^{hi}$CD69$^{+}$CD24$^{hi}$ (middle panel) and mature TCR$\beta^{hi}$CD69$^{-}$CD24$^{lo}$ T cell populations (bottom panel) in the thymus of mice with indicated genotypes. Data is representative of >3 experiments with 5 animals/group/experiment. **b**, FACS contour data plots showing T cell populations among TCRβ+ T cells in the spleen/LN of CD4+/+ and *Cd4*$^{E4a\Delta/\Delta}$ mice. Data is representative of >3 experiments with 5 animals/group/experiment. **c**, Bar graph quantifying CD4 MFI on different CD4 T cell populations from the thymus from mice with the indicated genotypes. Data are expressed as mean ± SEM of individual mice (n = 5 of two independent experiments). ****p<0.0001 (2-way ANOVA and Bonferroni test). **d**, Bar graph quantifying CD4 MFI on TCRβ+ CD4 T cell populations from the spleen/LN of mice with the indicated genotypes. Data are expressed as mean ± SEM of individual mice (n = 5 of two independent experiments). ***p<0.001, ****p<0.0001 (One-way ANOVA and Bonferroni test)

a

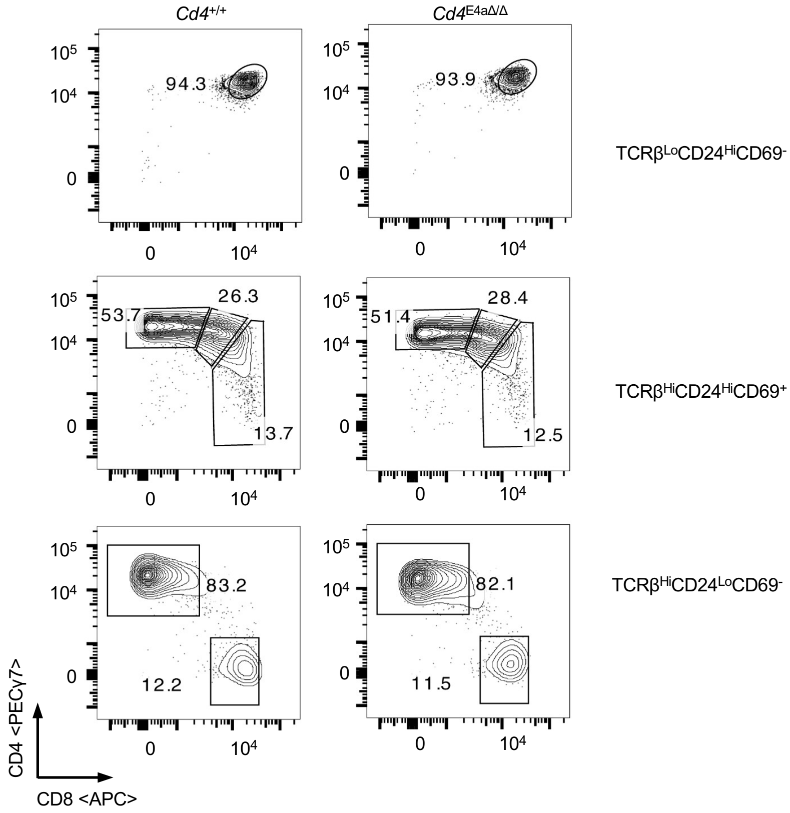


b

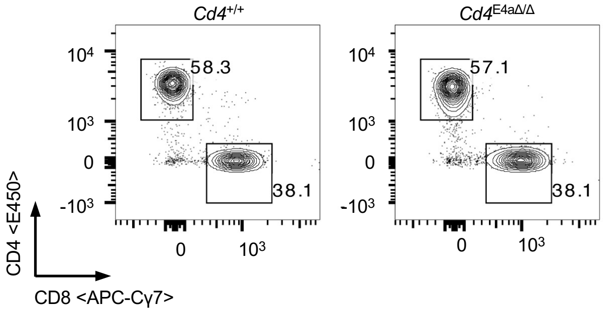


c

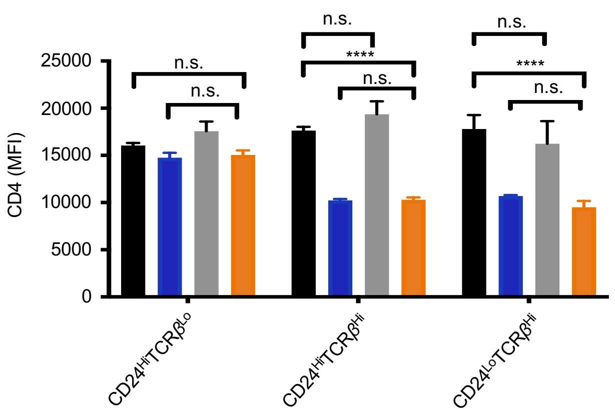


d

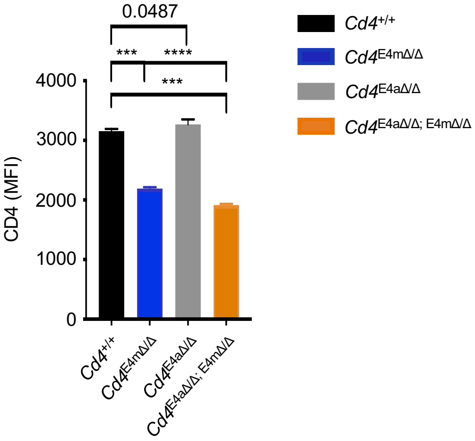


Supplementary Figure 2

**Supplementary Fig. 3: Lack of DNA demethylation during development affects the function of E4m/E4a in effector CD4-lineage T cells**. **a**, Heatmap (Top) and IGV snapshot (bottom) of the *Cd4* locus depicting individual CpGs with MM9 coordinates captured by CATCH-Seq in regions proximal to E4p (green) and E4a (blue). Differential CpGs flanking E4a are highlighted in red boxes. **b**, Heatmap depicting percent CpG methylation in control $CD4^+$ (Tet1/$3^{flox/flox}$ ), Tet1/$3^{cDKO}$ $CD4^+$ mature thymocytes and Tet1/$3^{cDKO}$ $CD4^+$ naïve peripheral T cells for CpGs from +6200 to −669 relative to the *Cd4* TSS (Chr6:124832027–124838896; mm9). A red line underlines CpGs in E4m (indicated by the gap in the mutant mice) and a black arrow indicates the *Cd4* TSS. CATCH-seq was performed on genomic DNA from sorted populations of $TCR\beta^{hi}CD24^{lo}CD69^{-}CD4^{+}CD8^{-}$ thymocytes or $CD4^{+}TCR\beta^{+}CD62L^{hi}$ $CD44^{-}$ T cells from LN/Spleen. Replicates are from 2 independent mice. **c**, Histone H3K4me3 modifications assessed by ChIP-qPCR in sorted naïve CD4 T cells activated *in vitro* for 120hrs. Data are averaged from two biological replicates/genotype and representative of 2 experiments. *p<0.05 (Student t tests). **d**, Histone H3K9me3 modifications assessed by ChIP-qPCR in sorted naïve CD4 T cells activated *in vitro* for 120hrs. Data are averaged from two biological replicates/genotype and representative of 2 experiments. *p<0.05 (Student t tests). **e**, Bar graph quantifying CD4 gMFI on T cells from indicated genotypes treated with DMSO or A-485 and analyzed 72hrs post activation with anti-CD3/CD28 *in vitro*. Cells were treated with DMSO vehicle control or indicated doses of A-485 18 hrs post-activation. Data shown is representative of 3 experiments and expressed as mean ± SD of 4 technical replicates from 2 independent mice/group. ****p<0.0001 (2-way ANOVA and Tukey’s multiple comparison test)

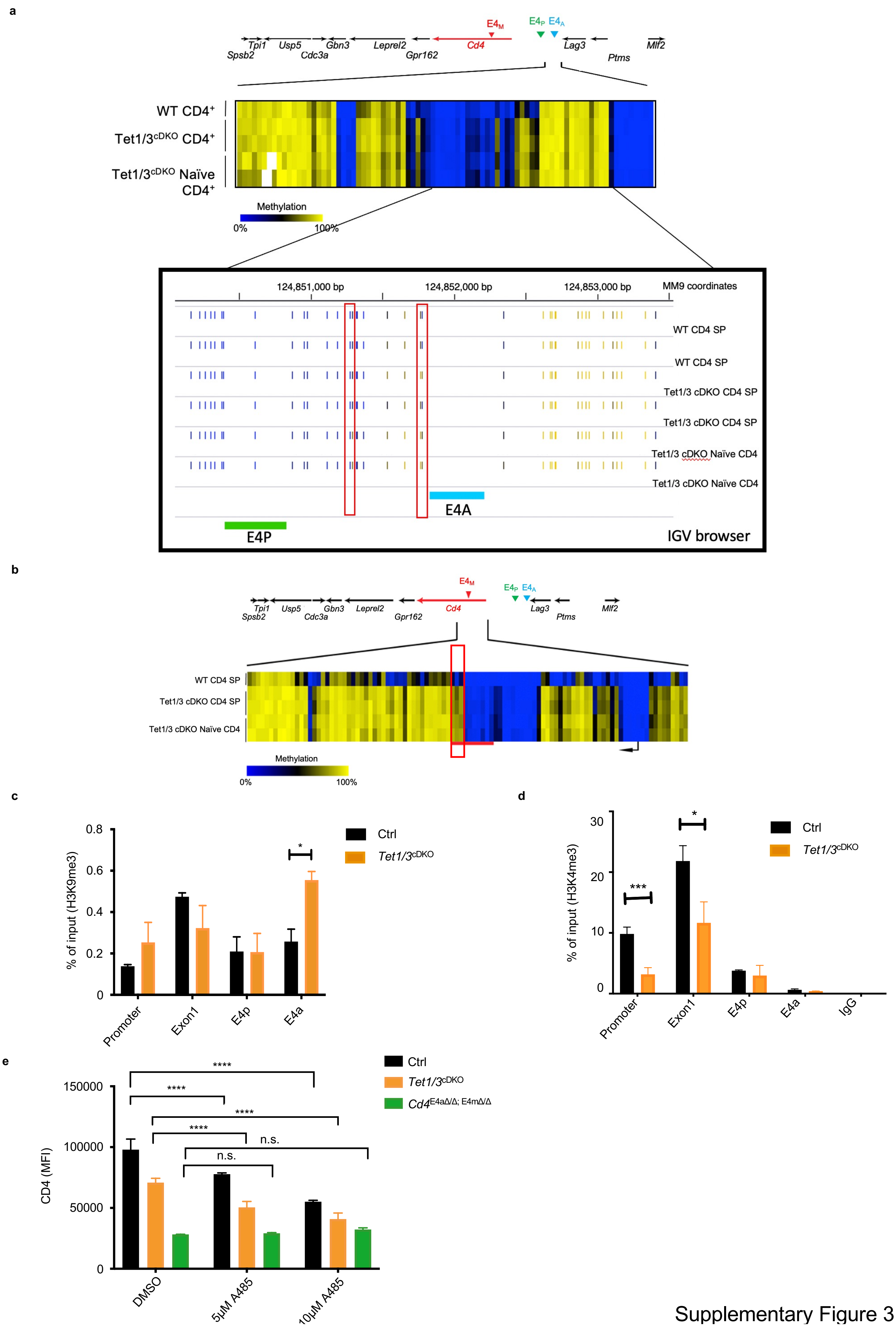


Supplementary Figure 3

**Supplementary Fig. 4: Reduced enhancer activity as a result of DNA methylation leads to *Cd4* promoter silencing during replication of effector CD4$^+$ T cells. a,** FACS plot showing loss of CD4 expression 96hrs after *in vitro* proliferation of activated CD4$^+$ T cells isolated from RorcCre$^{Tg}$ *Cd4*$^{S4fl/fl}$ and RorcCre$^{Tg}$ *Cd4*$^{S4fl/fl}$ *Dnmt3a*$^{fl/fl}$ mice. Data is representative of 3 experiments with 2 animals/group/experiment. **b,** Bar graph quantifying the % of CD4 T cells from indicated genotypes losing CD4 expression 96hrs after *in vitro* proliferation. Data is a summary of 3 independent experiments. **p<0.001 (Student t test)**. c,** Bar graph quantifying CD4 gMFI 96hrs post *in vitro* activation. Data shown is representative of 3 experiments and expressed as mean ± SD of 3 technical replicates. Significant p values are indicated (Student t-test)**. d,** Bar graph quantifying CD4 gMFI in activated T cells from RorcCre$^{Tg}$ *Cd4*$^{S4fl/fl}$ *Dnmt3a*$^{fl/fl}$ mice, 96hrs post transduction with an shRNA against Renilla (control), Dnmt1 or Dnmt3a. Data shown is representative of 2 experiments and expressed as mean ± SD of 2 technical replicates from 2 biological replicates. *p<0.05 (One-way ANOVA and Sidak's multiple comparison test)**. e,** Model depicting the importance of E4m and E4a enhancer activities and CD4 gene expression in resting CD4+ thymic T cells versus activated effector CD4 T cells. E4a and E4m enhancer activities are critical for maintaining H3K4me3 levels at the promoter during replication. A lack of enhancer activity (E4m/E4m doubly deficient T cells) or reduced E4a/E4m enhancer activities (Tet1/3-deficient T cells) leads to a gain of *de novo* methylation and suppression of promoter activity and ultimately the loss of CD4 gene expression.

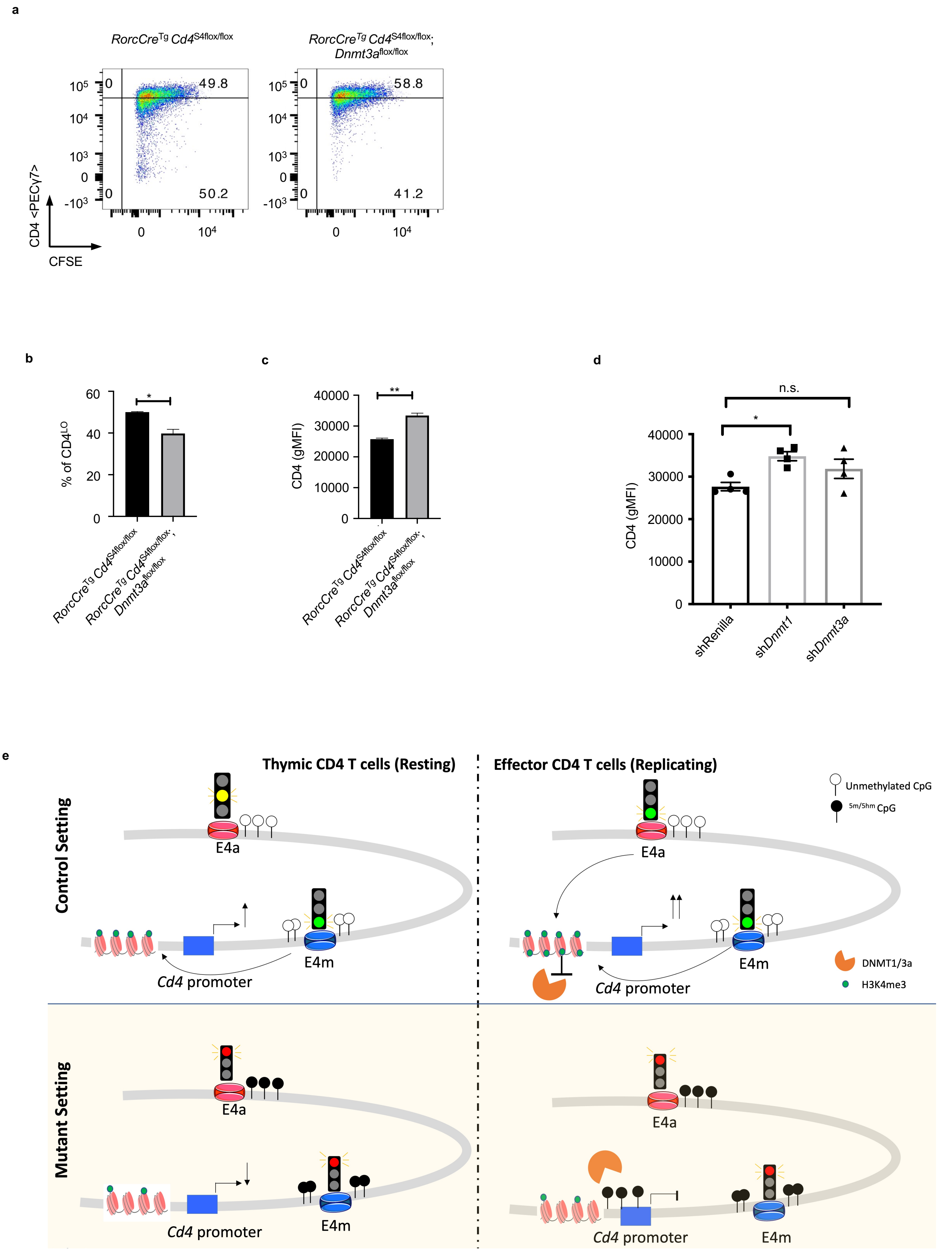


Supplementary Figure 4

**Supplementary Fig. 5: Reduced CD4 expression in effector CD4 T cells impairs parasitic clearance during Leishmaniasis. a,** FACS contour plots showing CD4 expression on cells gated on $CD11a^{+}CD44^{hi}$ T cells among $CD8^{-}$ $TCR\beta^{+}$ T cells. Cells were isolated from the dLNs of Leishmania infected mice at day 9 post infection. **b, c**, quantification of CD4 MFI on $CD11a^{+}CD44^{hi}$ T cells among $CD8^{-}$ $TCR\beta^{+}$ T cells from d9 infected mice with indicated genotypes.

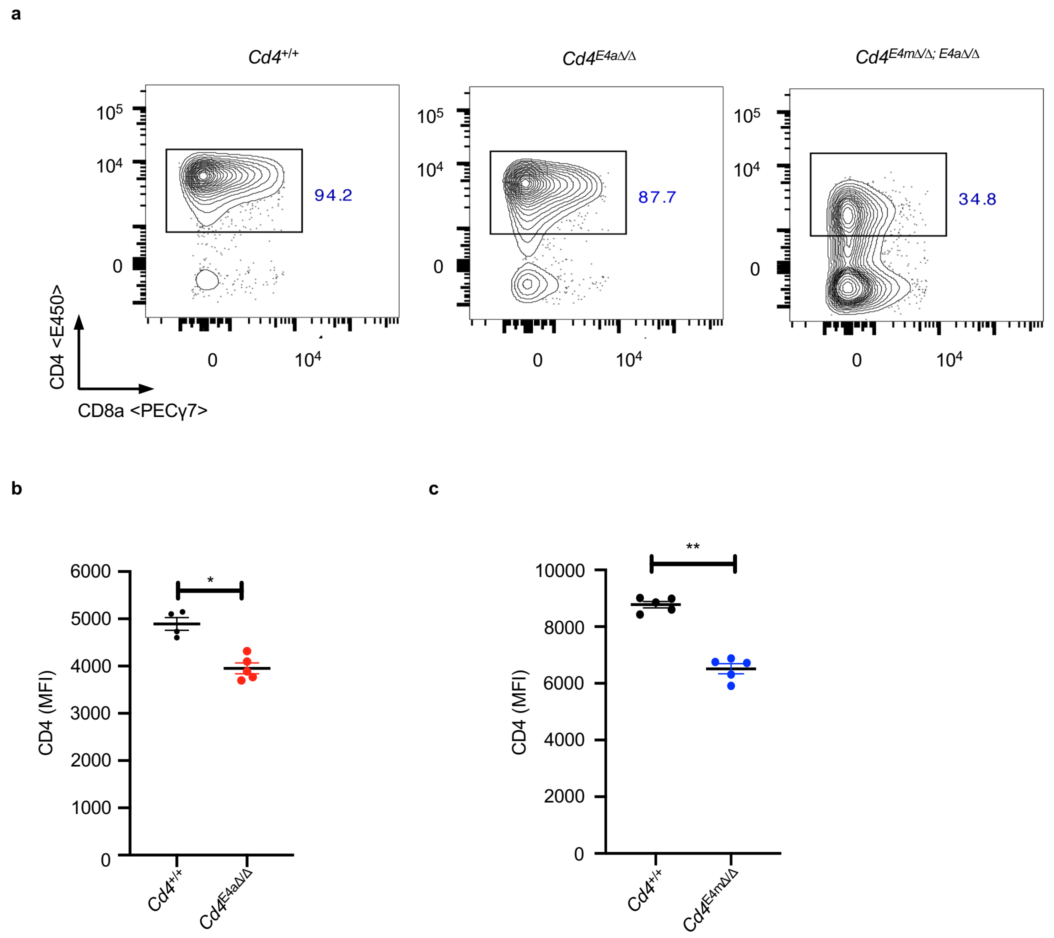


Supplementary Figure 5

**Supplementary Fig. 6: Reduced CD4 expression impairs the differentiation of Th1 cells during Leishmaniasis. a,** FACS contour plot depicting proportions of $CD11a^{+}CD44^{hi}$ T cells among $CD8^{-}$ $TCR\beta^{+}$ T cells (Top panel) or $CD11a^{+}CD49d^{hi}$ T cells among $CD8^{-}$ $TCR\beta^{+}$ T cells in the draining inguinal LNs of *L.major*-infected mice, 9 days post infection. Data is representative of more than 3 independent experiments. **b,** Absolute number of $CD8^{-}$ $TCR\beta^{+}$ T cells from the draining inguinal LNs of d28 *L.major*-infected mice. **c,** MFI of IFNγ+ cells among $CD8^{-}$ $TCR\beta^{+}$ $CD11a^{+}CD44^{hi}$T cells from the draining inguinal LNs of d28 *L.major*-infected mice or uninfected mice after restimulation overnight with soluble leishmania antigen *in vitro*. Data shown is representative of 2 independent experiments with 4-5 mice/group with technical replicates. *p<0.05, **p<0.01 (One-way ANOVA and Kruskal- Wallis test)**. d,** IL-4 levels in the supernatants from homogenized footpads of *L. major*–infected mice (5-6 mice/group) on day 28 analyzed by multiplex ELISA (Mann-Whitney test)

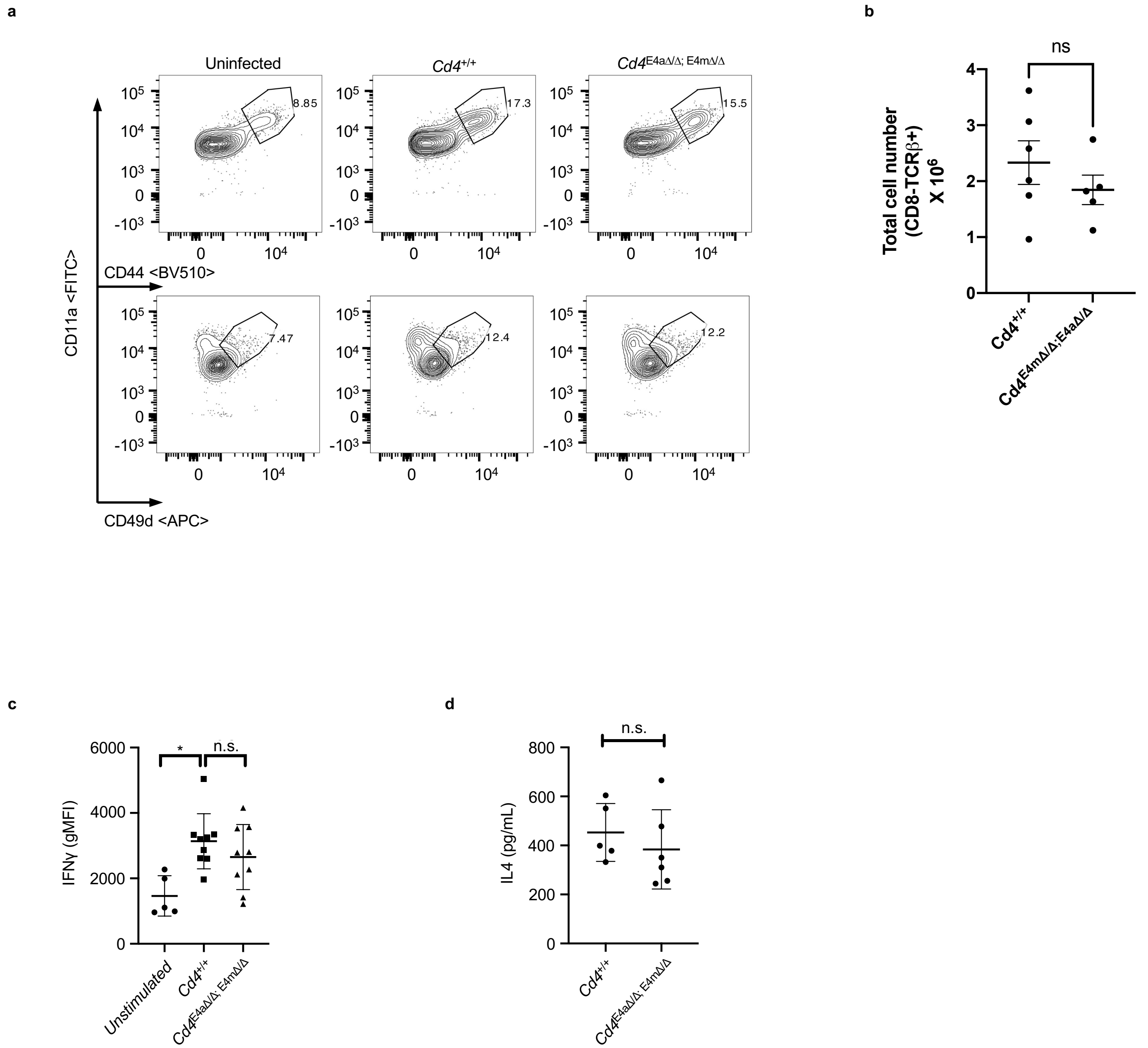


Supplementary Figure 6

**Supplementary Fig. 7: TET-mediated demethylation during thymic development is critical for optimal gene function in effector T cells a,** Scatter plot showing a fold change >2 in gene expression from DN3 to CD4+ thymic T cell differentiation (y-axis) versus presence of 5hmC in CD4+ T cells (log2 CMS-IP/Input >2). Number of genes in each quadrant is shown and depicted genes are listed in Supplementary Table 1. **b,** Scatter plot depicting change in gene expression from DN3 to CD4+ thymic T cell differentiation ((log2FoldChange) > 1) versus change in intragenic 5hmC in CD4+ compared to CD4+CD8+ DP thymic T cells. Genes with intragenic 5hmC (log2 CMS-IP/Input > 2) in at least one of the two cell types were considered and genes with an absolute difference of >0.5 between them were plotted. Number of genes in each quadrant is shown and depicted genes are listed in Supplementary Table 2. **c,** Venn diagram showing the number of genes which display novel chromatin accessibility peaks upon TCR activation among the group of 350 genes that are upregulated during CD4SP differentiation in the thymus (a fold change >2 in gene expression from DN3 to $CD4^{+}$SP ) and undergo DNA demethylation (Intragenic 5hmC log2 CMS-IP/Input >2). **d,** Representative picture showcasing the lack of splenomegaly and lympho-adenopathy in Rorc(t)$Cre^{Tg}$ $Tet1^{fl/fl}$ $Tet2^{fl/+}$ $Tet3^{fl/fl}$ mice (Top panel) versus severe splenomegaly and lympho-adenopathy in 5 weeks old Rorc(t)$Cre^{Tg}$ $Tet1^{fl/fl}$ $Tet2^{fl/fl}$ $Tet3^{fl/fl}$ mice (Bottom). **e,** Representative FACS contour plots depicting proportions of CD4 and CD8 T cells among $TCR\beta^{+}$ T cells (Top panel) and proportions of naïve $CD4^{+}$ $CD62L^{hi}$ $CD44^{-}$ T cells among $CD4^{+}$ $TCR\beta^{+}$ T cells (Bottom Panel) in the spleen and LN of 5-6 weeks old control and Rorc(t)$Cre^{Tg}$ $Tet1^{fl/fl}$ $Tet2^{fl/+}$ $Tet3^{fl/fl}$ mice. **f ,** Histogram overlay of CD4, ThPOK, CD5 and CD6 expression on peripheral $CD4^{+}$ $TCR\beta^{+}$ T cells from the LN of Rorc(t)$Cre^{Tg}$ $Tet1^{fl/fl}$ $Tet2^{fl/+}$ $Tet3^{fl/fl}$ mice (**top**) and activated $CD4^{+}$ $TCR\beta^{+}$ T cells FACS-sorted from the LN of Rorc(t)$Cre^{Tg}$ $Tet1^{fl/fl}$

$Tet2^{fl/+}$ $Tet3^{fl/fl}$ mice and activated *in vitro* for 72hrs with anti-CD3/CD28 (bottom). Data is representative of >3 independent experiments. **g**, % of cells with hi or low gene expression per cell division cycle following in vitro activation for 72hrs with anti-CD3/CD28. **h,** Normalized CD4 MFI expression in CD4 T cells from $Rag^{-/-}$ mice at day 7 post transfer. Naïve FACS-sorted WT CD45.1 and Rorc(t)$Cre^{Tg}$ $Tet1^{fl/fl}$ $Tet2^{fl/+}$ $Tet3^{fl/fl}$ mice CD45.2 T cells were transferred at a 1:1 ratio and transferred into $Rag^{-/-}$. Expression of the respective protein was normalized to the WT control. Data is representative of 2 independent experiments with 3-5 mice/experiment.

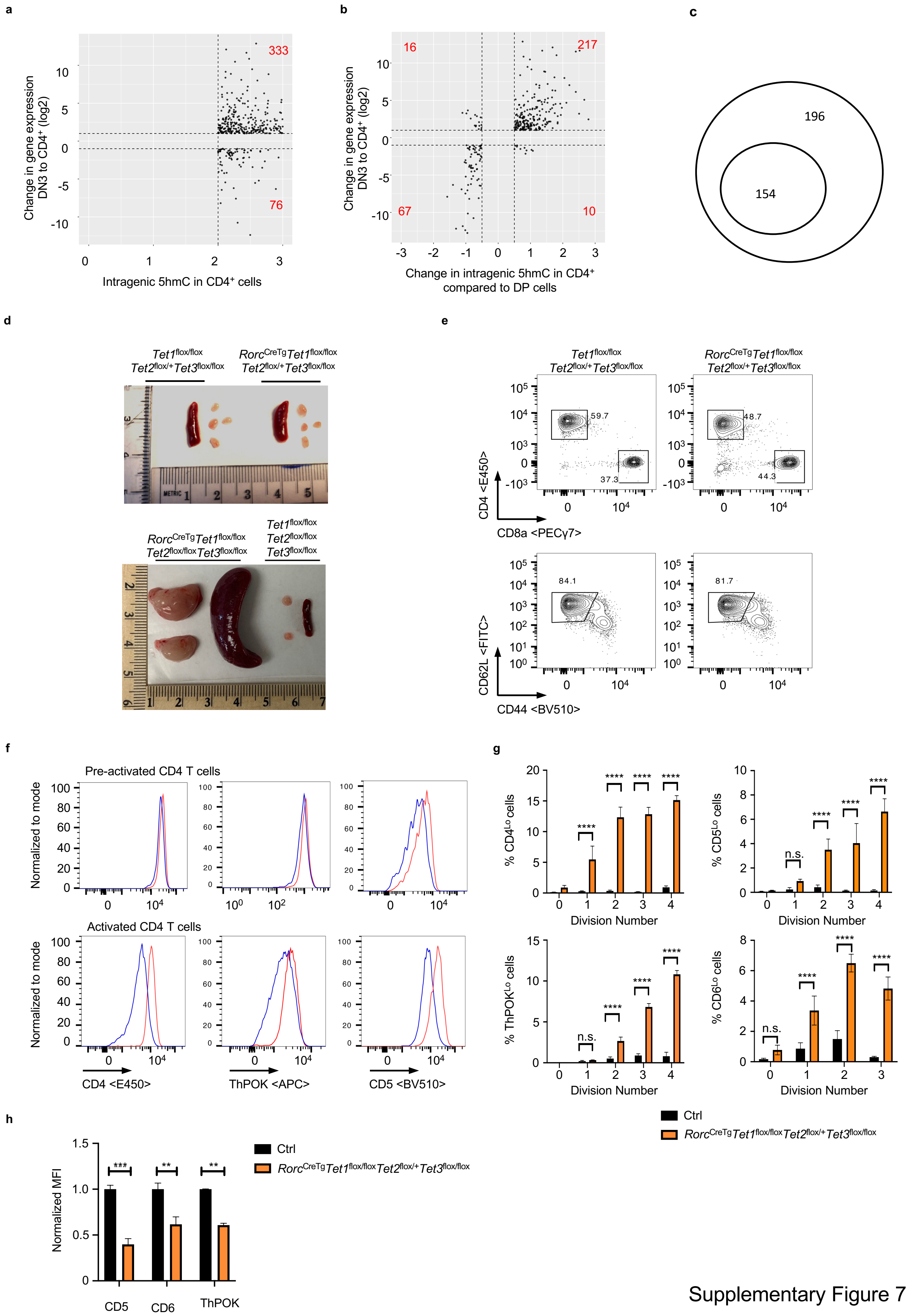


Supplementary Figure 7
